# Spatial control of nucleoporin assembly by Fragile X-related proteins

**DOI:** 10.1101/767202

**Authors:** Arantxa Agote-Arán, Stephane Schmucker, Katerina Jerabkova, Inès Jmel Boyer, Alessandro Berto, Laura Pacini, Paolo Ronchi, Charlotte Kleiss, Laurent Guerard, Yannick Schwab, Hervé Moine, Jean-Louis Mandel, Sebastien Jacquemont, Claudia Bagni, Izabela Sumara

## Abstract

Nucleoporins (Nups) build highly organized Nuclear Pore Complexes (NPCs) at the nuclear envelope (NE). Several Nups assemble into a sieve-like hydrogel within the central channel of the NPCs to regulate nucleocytoplasmic exchange. In the cytoplasm, a large excess of soluble Nups has been reported, but how their assembly is restricted to the NE is currently unknown. Here we show that Fragile X-related protein 1 (FXR1) can interact with several Nups and facilitate their localization to the NE during interphase through a microtubule and dynein-dependent mechanism. Downregulation of FXR1 or closely related orthologs FXR2 and Fragile X mental retardation protein (FMRP) leads to the accumulation of cytoplasmic Nup protein condensates. Likewise, several models of Fragile X syndrome (FXS), characterized by a loss of FMRP, also accumulate cytoplasmic Nup aggregates. These aggregate-containing cells display aberrant nuclear morphology and a delay in G1 cell cycle progression. Our results reveal an unexpected role for the FXR protein family and dynein in the spatial regulation of nucleoporin assembly.

**Highlights:** Cytoplasmic nucleoporins are assembled by Fragile X-related (FXR) proteins and dynein

FXR-Dynein pathway downregulation induces aberrant cytoplasmic aggregation of nucleoporins

Cellular models of Fragile X syndrome accumulate aberrant cytoplasmic nucleoporin aggregates.

FXR-Dynein pathway regulates nuclear morphology and G1 cell cycle progression

**eTOC Blurb:** Nucleoporins (Nups) form Nuclear Pore Complexes (NPCs) at the nuclear envelope. Agote-Arán at al. show how cells inhibit aberrant assembly of Nups in the cytoplasm and identify Fragile X-related (FXR) proteins and dynein that facilitate localization of Nups to the nuclear envelope and control G1 cell cycle progression.

Graphical abstract

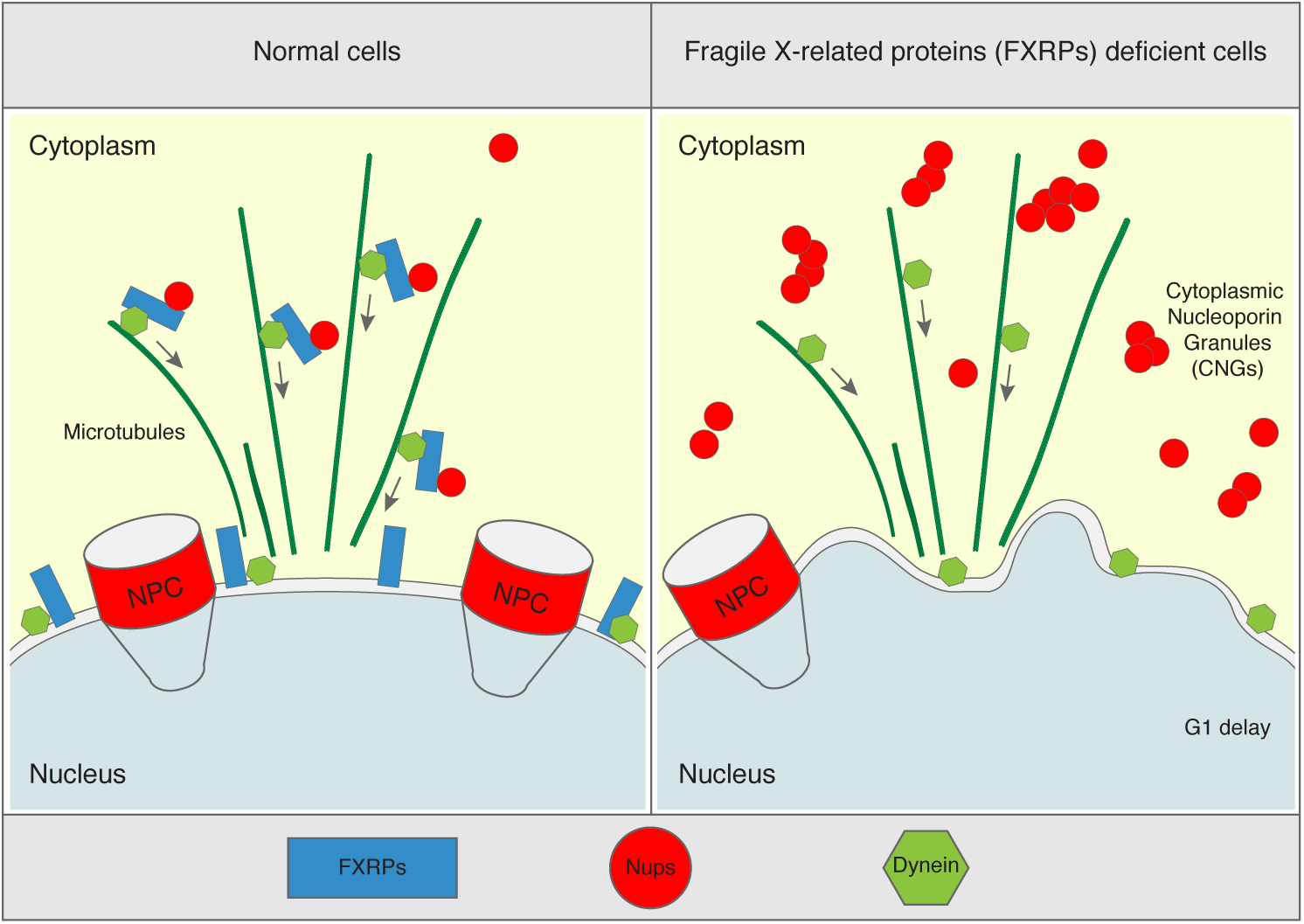

## Introduction

Formation of supramolecular assemblies and membrane-less organelles such as the nucleolus, Cajal bodies, nuclear speckles, stress granules (SG), P-bodies, germ granules and PML bodies are important for cellular homeostasis (Boeynaems et al., 2018). Among the factors controlling their formation and turnover is the presence of intrinsically disordered regions (IDRs) in protein components, their ability to form multivalent protein-protein, and protein-RNA interactions (Feng et al., 2019) and proteins’ local concentration. Indeed, many RNA-binding proteins (RBPs) have the ability to demix into liquid states (liquid droplets), which can be subsequently transformed into pathological amyloids both *in vitro* and in cells (Harrison and Shorter, 2017; Lin et al., 2015; Shorter, 2019). Formation and clearance of pathological protein condensates have been linked to many neurological disorders (Shin and Brangwynne, 2017). One example of a large protein assembly consisting of IDR-containing proteins is the Nuclear Pore Complex (NPC), which plays an essential role in cellular homeostasis (Knockenhauer and Schwartz, 2016; Sakuma and D’Angelo, 2017).

NPCs are large, multisubunit protein complexes (Beck and Hurt, 2017a; Hampoelz et al., 2019) spanning the nuclear envelope (NE) that constitute the transport channels controlling the exchange of proteins and mRNAs between the nucleus and the cytoplasm. They are built from roughly 30 different nucleoporins (Nups) each present in multiple copies in the NPCs. The ring-like NPC scaffold is embedded in the NE and shows highly organized eight-fold symmetry (Knockenhauer and Schwartz, 2016).

In contrast, the central channel of the NPC is formed from Nups containing disordered elements characterized by the presence of phenylalanine-glycine (FG) repeats, the so-called FG-Nups. The FG-Nups have the ability to phase separate into sieve-like hydrogels that constitute a selective and permeable barrier for diffusing molecules and transported cargos through the NPCs (Schmidt and Görlich, 2016). This ability of the FG-Nups to form permeable amyloid-like hydrogels can also be reconstituted *in vitro* and is highly conserved through the evolution (Frey and Görlich, 2007; Frey et al., 2006; Schmidt and Görlich, 2015). The cohesive abilities of FG-Nups allow not only for the formation of the permeability barrier but also for building the links with the structural scaffold elements of the NPC (Onischenko et al., 2017). The non-FG-Nups can also form condensates in cells as they are sequestered in the stress granules (Zhang et al., 2018) and in various pathological aggregates in the nucleus and in the cytoplasm (Hutten and Dormann, 2019; Li and Lagier-Tourenne, 2018). A fraction of cytoplasmic nucleoporins was also identified in the promyelocytic leukemia protein (PML)-positive structures, the so-called CyPNs (cytoplasmic accumulations of PML and nucleoporins), which could move on microtubules to dock at the NE (Jul-Larsen et al., 2009), although the cellular roles of the CyPNs remain to be understood. This indicates that Nups have an intrinsic capacity to aberrantly assemble, suggesting protective mechanisms may exist to prevent it in the cell. Indeed, Nups are synthesized as soluble proteins in the cytoplasm and in Drosophila embryos a large excess of soluble Nups has been reported (Onischenko et al., 2004). How the balance of soluble Nups is controlled, and what factors regulate the localized assembly of Nups is currently unknown.

The Fragile X related (FXR) proteins (FXR1, FXR2 and Fragile X mental retardation protein (FMRP)) are a family of RNA-binding proteins displaying a high degree of sequence and structural similarity (Li and Zhao, 2014). Silencing of the *FMR1* gene that encodes the FMRP protein (Santoro et al., 2012) leads to Fragile X syndrome (FXS), the most common form of inherited intellectual human disability worldwide, for which no efficient therapy exists to date (Mullard, 2015). Here we identify an unexpected role for the FXR protein family and dynein in the spatial regulation of nucleoporin assembly.

## Results

### FXR1 protein localizes to the NE and interacts with Nups

FXR1 co-localizes with various cytoplasmic protein-RNA assemblies, but it is also present in the nuclear compartment in human cells (Oldenburg et al., 2014; Tamanini et al., 1999). In search for possible cellular functions of FXR1 independent of its role in RNA binding, we performed immunoprecipitations (IPs) of stably expressed GFP-FXR1 protein and analysed the interacting partners by mass spectrometry. Of the interacting proteins, including the known FXR1 partners, FXR2 and FMRP, four nucleoporins (Nups), gp210, Nup188, Nup133 and Nup85, were detected specifically in GFP-FXR1 but not in GFP-control IPs (Table 1). We confirmed the GFP-FXR1 interaction with endogenous Nup133 and Nup85, which are components of the evolutionary conserved Nup107-160 NPC sub-complex also called the Y-complex (Figure 1A) (Beck and Hurt, 2017b; Knockenhauer and Schwartz, 2016). IP of stably expressed GFP-Nup85 also demonstrated an interaction with endogenous FXR1 in HeLa cells (Figure 1B), and both Nup85 and Nup133 immunoprecipitated endogenous FXR1 in HEK293T cells (Figure 1C).

**Figure 1.**
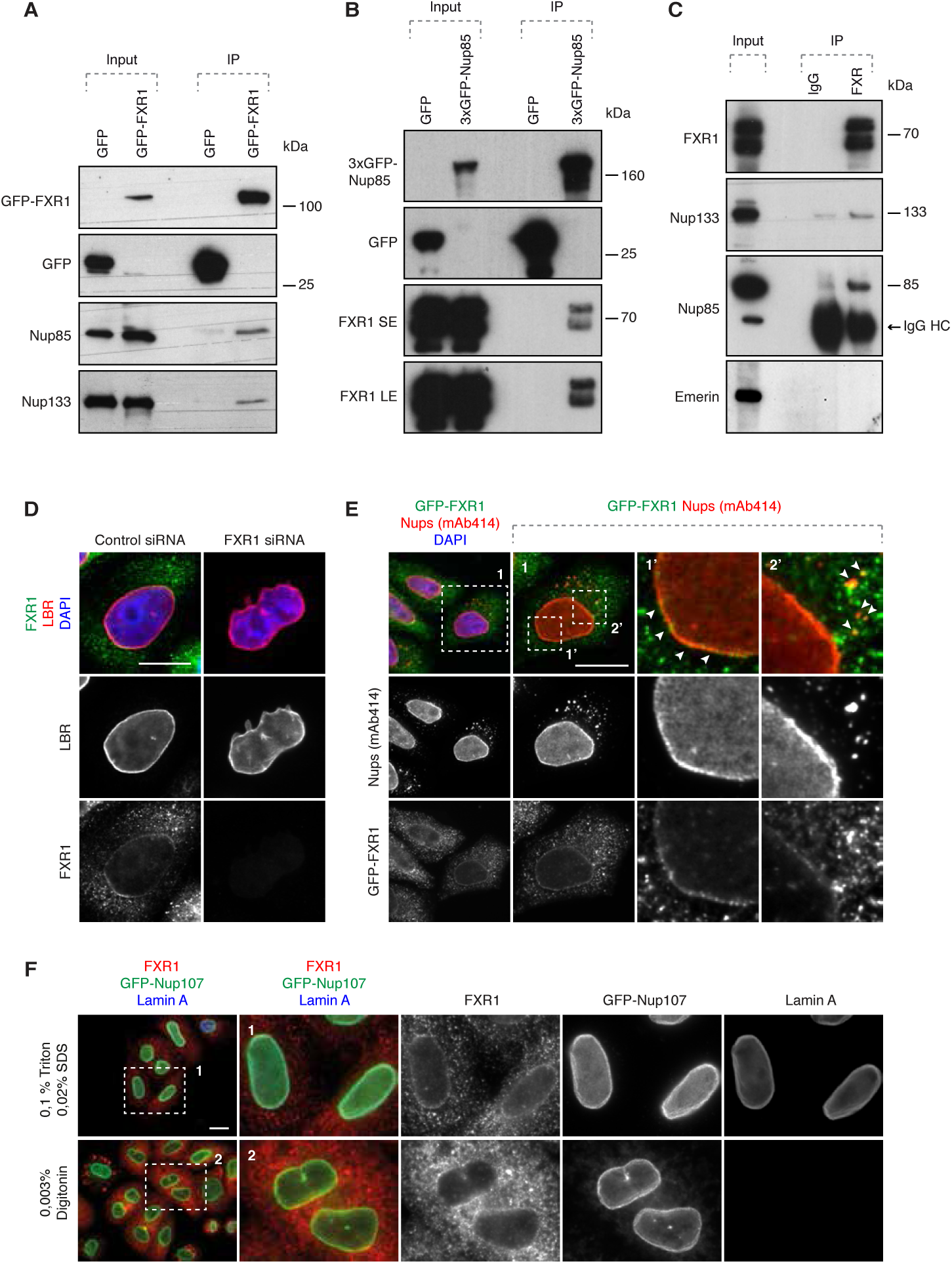
FXR1 protein localizes to the NE and interacts with NUPs. **(A)** Lysates of HeLa cells stably expressing GFP alone or GFP-FXR1 were subjected to immunoprecipitation using GFP-Trap beads (GFP-IP) and analysed by Western blot. **(B)** Lysates of HeLa cells stably expressing GFP alone or 3xGFP-Nup85 were immunoprecipitated using GFP-Trap beads (GFP-IP) and analysed by Western blot (SE, short exposure, LE, long exposure). **(C)** Immunoprecipitation from HEK293T cell lysates using FXR1 antibody or IgG analysed by Western blotting. The arrow points to the heavy chain of IgG (IgG HC). **(D)** HeLa cells were treated with indicated siRNAs, synchronized by double thymidine block, and released for 12 hours and analysed by immunofluorescence microscopy for the lamin B receptor (LBR) to mark the NE, and FXR1. **(E)** HeLa cells stably expressing GFP-FXR1 were analysed by immunofluorescence microscopy for GFP and mAb414, which binds FG-Nups. The magnified framed regions are shown in the corresponding numbered panels. The arrowheads indicate NE and cytoplasmic localization of GFP-FXR1. **(F)** HeLa cells stably expressing GFP-Nup107 were synchronized by double thymidine block and released for 12 hours, permeabilized with Triton/SDS or digitonin for antibodies to access the nucleus, and analysed by immunofluorescence microscopy. Bars are 5 µm.

**Table 1:** Mass spectrometry analysis of GFP-FXR1-interacting partners. HeLa cells expressing GFP alone or GFP-FXR1 were immunoprecipitated using agarose GFP-Trap A beads (GFP-IP) and analysed by mass spectrometry. For each hit, the uniprot accession number, percentage of peptide coverage, number of unique identified peptides (3 or more), number of amino acids in the protein sequence, predicted molecular weight and PI of proteins are depicted.

| Accession | Description | ΣCoverage | # Unique Peptides | MW [kDa] | calc. pI |
| --- | --- | --- | --- | --- | --- |
| P51114-2 | Isoform 2 of Fragile X mental retardation syndrome-related protein 1 OS=Homo sapiens GN=FXR1 - [FXR1_HUMAN] | 70,50 | 41 | 60,8 | 6,39 |
| P51114 | Fragile X mental retardation syndrome-related protein 1 OS=Homo sapiens GN=FXR1 PE=1 SV=3 - [FXR1_HUMAN] | 64,41 | 40 | 69,7 | 6,15 |
| Q14204 | Cytoplasmic dynein 1 heavy chain 1 OS=Homo sapiens GN=DYNC1H1 PE=1 SV=5 - [DYHC1_HUMAN] | 26,39 | 19 | 532,1 | 6,40 |
| P51116 | Fragile X mental retardation syndrome-related protein 2 OS=Homo sapiens GN=FXR2 PE=1 SV=2 - [FXR2_HUMAN] | 39,38 | 19 | 74,2 | 6,23 |
| Q06787-6 | Isoform 5 of Fragile X mental retardation protein 1 OS=Homo sapiens GN=FMR1 - [FMR1_HUMAN] | 45,76 | 18 | 66,4 | 8,38 |
| Q8TEM1 | Nuclear pore membrane glycoprotein 210 OS=Homo sapiens GN=NUP210 PE=1 SV=3 - [PO210_HUMAN] | 8,85 | 4 | 205,0 | 6,81 |
| Q5SRE5-2 | Isoform 2 of Nucleoporin NUP188 homolog OS=Homo sapiens GN=NUP188 - [NU188_HUMAN] | 3,72 | 4 | 182,2 | 6,83 |
| Q8WUM0 | Nuclear pore complex protein Nup133 OS=Homo sapiens GN=NUP133 PE=1 SV=2 - [NU133_HUMAN] | 14,10 | 3 | 128,9 | 5,10 |
| Q9BW27-2 | Isoform 2 of Nuclear pore complex protein Nup85 OS=Homo sapiens GN=NUP85 - [NUP85_HUMAN] | 12,99 | 3 | 52,6 | 6,52 |

Both endogenous FXR1 and GFP-FXR1 localized to the nuclear envelope (Figure 1D, E) and also occasionally to small cytoplasmic foci labelled by the monoclonal antibody mAb414, which recognizes a panel of phenylalanine-glycine (FG) repeat-containing Nups (FG-Nups) (Fig. 1E). FXR1 localization to the NE and the cytoplasmic vesicles was abolished by treatment with FXR1 siRNA (Figure 1D), demonstrating antibody specificity, and digitonin treatment, which selectively permeabilizes the plasma membrane while leaving the NE intact, revealed that FXR1 was largely restricted to the outer nuclear membrane (ONM) (Figure 1F). We conclude that FXR1 interacts with Nups, and localizes both to the ONM and to cytoplasmic foci containing Nups.

### FXR1 inhibits aberrant assembly of cytoplasmic Nups

To assess the biological function of the FXR1-Nup interactions, we treated cultured human cells with FXR1-specific siRNA oligonucleotides. Downregulation of FXR1 led to a dramatic accumulation of FG-Nups in the cytoplasm in the form of irregular aggregate-like assemblies of various sizes (Figure 2A, B and Figure S1, S3A). These Nup assemblies were observed using two different siRNAs targeting FXR1 and could be largely rescued by stable ectopic expression of a form of GFP-FXR1 that is resistant to one of the siRNAs used (Figure 2B and Figure S1). Defects in Nup localization in the FXR1-deficient cells was likely not due to defective assembly of the nuclear lamina, which is a key structural component of the inner nuclear membrane, as downregulation of FXR1 did not affect recruitment of the lamin B receptor (LBR) (Figure 1D), lamin A (Figure S2A, B), lamin B1 (Figure S2C, D), emerin or Lap2β (Figure S2E-G) to the NE in interphase or telophase cells. Downregulation of FXR1 led to the cytoplasmic retention and co-localization in aggregates of at least 10 Nups spanning several functional and structural NPC groups, including FG-Nups (Nup98, Nup214; and RanBP2); transmembrane Nups (Gp210 and POM121); Y-complex Nups (Nup133 and Nup85, stably expressed GFP-Nup85 and stably expressed GFP-Nup107) (Figure 2C, Figure S3B-F, S4A, B, S5D, S7A) as well as Nup88 and NPC-associated the RanGTPase activating protein (RanGAP1). Absent from the Nup aggregates were the nuclear ring Nup ELYS and the inner nuclear basket component Nup153, although their levels at the NE were both reduced (Figure 2C, Figure S4D, S5A-C). FXR1 downregulation also reduced the NE localization of FG-Nups, RanBP2 and stably expressed GFP-Nup107 (Figure 2D-F). Collectively, these data show that loss of FXR1 interferes with Nup localization to the NE, and instead induces their inappropriate assembly in the cytoplasm.

**Figure 2.**
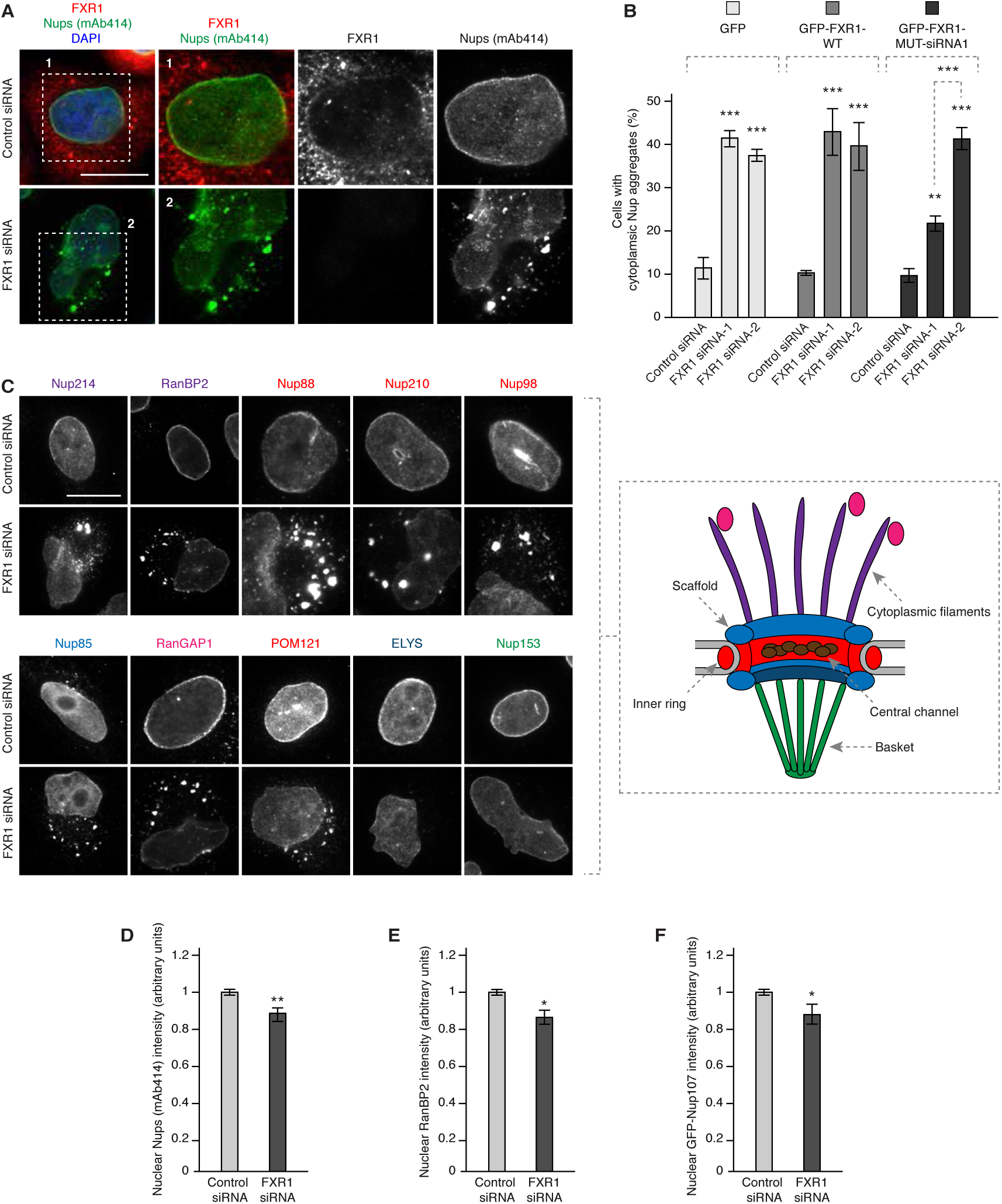
FXR1 depletion induces aberrant assembly of cytoplasmic Nups. **(A)** HeLa cells were treated with the indicated siRNAs, synchronized by double thymidine block and release for 12 hours and analysed by immunofluorescence microscopy. The magnified framed regions are shown in the corresponding numbered panels. Additional representative images of cells are shown in Figure S3A. **(B)** HeLa cells stably expressing GFP, GFP-FXR1 wild type (WT) and GFP-FXR1 mutated in the sequence recognized by FXR1 siRNA-1 (GFP-FXR1-MUT-siRNA1) were treated with the indicated siRNAs, synchronized by double thymidine block, released for 24 hours and then analysed by immunofluorescence microscopy. The percentage of cells with cytoplasmic nucleoporin aggregates (n=1000) was quantified. The corresponding representative pictures are shown in Figure S1 and the corresponding Western blot analysis is shown in Figure 6B. **(C-F)** HeLa cells were treated with the indicated siRNAs, synchronized by double thymidine block, released for 12 hours and analysed by immunofluorescence microscopy. Nups present in different NPC subcomplexes are depicted in the color code corresponding to the NPC scheme shown on the right. Additional or complementary representative images and channels of cells depicted in (C) are shown in Figure S3 B-F, S4 B-D and S5B-D. Nuclear intensity of FG-Nups labelled by mAb414 (D), RanBP2 (E) and GFP-Nup107 (F) was quantified (n=300). Bars are 5 µm.

### Loss of FXR1 affects NPC assembly during early interphase

To date, two temporally and mechanistically distinct pathways of NPC assembly at the NE have been described during the cell cycle in higher eukaryotic cells (Weberruss and Antonin, 2016). In the post-mitotic pathway, ELYS initiates NPC assembly on segregated chromosomes, while during interphase, both Nup153 and POM121 drive *de novo* assembly of NPCs into an enclosed NE (D’Angelo et al., 2006; Doucet et al., 2010; Vollmer et al., 2015). ELYS assembled normally on segregating chromosomes in anaphase and on decondensing chromatin in telophase in FXR1-deficient cells (Figure S5A). In addition, given we found that ELYS was not recruited to the cytoplasmic Nup aggregates whereas POM121 was (Figure 2C, Figure S5A, B, C), this suggests that loss of FXR1 affects Nup localization at the NE specifically during interphase. To corroborate this, we used live video microscopy of a reporter cell line stably expressing GFP-Nup85 and the chromatin marker histone H2B labelled with mCherry. FXR1 depletion did not affect progression and timing through mitosis or fidelity of chromosome segregation (Figure 3A, C). This was confirmed by analysis of the degradation of several mitotic factors in synchronized cells (Figure S6A, B). As expected, downregulation of FXR1 led to the accumulation of GFP-Nup85 in cytoplasmic aggregates (Figure 3A, B), with similar appearance and distribution to that observed in the fixed specimens, that co-localized with FG-Nups (Figure S4A), which was consistent with our earlier results analyzing endogenous Nup85. GFP-Nup85-positive aggregates became detectable in the cytoplasm on average 48 min after chromosome segregation and approximately 30 min after the onset of chromatin decondensation (Figure 3A, C). This indicates that loss of FXR1 affects Nup localization during early interphase. We considered that this effect may be mediated by a modulation of Nups levels. However, the protein levels of Nup85, Nup93, Nup133 and Nup155 (Figure S6A-C), and the mRNA levels of Nup85 and Nup133 (Figure S6D, E), as well as the known cell cycle-linked degradation of Nup85 (Figure S6A, B) and Nup133 dephosphorylation, which occurs during mitotic exit (Figure S6B), were unchanged upon depletion of FXR1. Together, our results suggest that loss of FXR1 affects Nup localization during early interphase.

**Figure 3.**
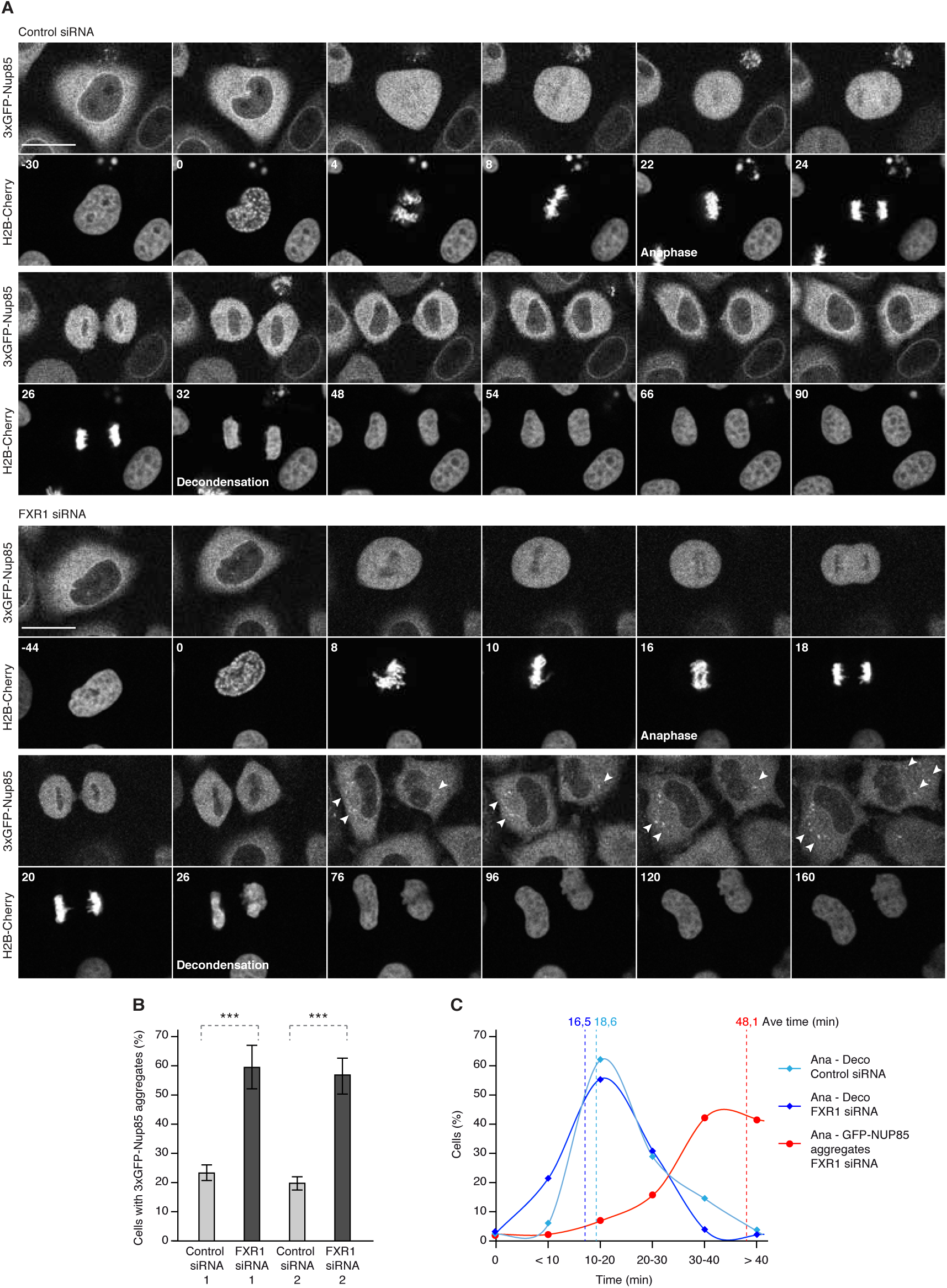
FXR1 regulates cytoplasmic Nups during early interphase. **(A-C)** HeLa cells stably expressing the chromatin marker histone H2B labelled with mCherry and 3xGFP-Nup85 were treated with indicated siRNAs, synchronized by double thymidine block, released and analysed by live video spinning disk confocal microscopy. The selected frames of both channels of the movies are depicted and time is shown in minutes. The onset of anaphase and chromatin decondensation are indicated. Arrowheads point to the cytoplasmic 3xGFP-NUP85 aggregates appearing during nuclear expansion of FXR1-deficient cells. Bars are 5 µm. The percentage of cells with cytoplasmic 3xGFP-Nup85 aggregates (n=73) was quantified in (B). **(C)** The distribution of percentage of cells that required the indicated times (min) for the transition from anaphase to DNA decondensation (Ana-Deco) in control (light blue) and FXR1 siRNA cells (dark blue), and of cells that required the indicated times (min) for the transition from anaphase to the appearance of the first detectable cytoplasmic 3xGFP-Nup85-positive structures in FXR1 siRNA cells (red). The average times are indicated by dotted lines in the corresponding color code.

### Nup aggregates are distinct from ALs, SGs and resistant to RNA degradation

The cytoplasmic Nup aggregates in FXR1-deficient cells could correspond to annulate lamellae (AL), which are preassembled NPCs embedded in the ER membrane (Merisko, 1989), as suggested by co-localization of various Nups in these assemblies. Indeed, cytoplasmic AL-NPCs can be inserted “en bloc” into an intact NE during embryogenesis in *Drosophila* (Hampoelz et al., 2016). A closer analysis by superresolution microscopy revealed an amorphous organization of the Nup aggregates in the perinuclear area of FXR1-deficient cells relative to the more regular, round shape of the small cytoplasmic Nup foci observed in the control cells (Figure S7A). Moreover, we were unable to detect any AL-typical structures, (characterized by parallel stacks of ER-membranes with embedded regularly spaced NPCs) in the FXR1-deficient cells by electron microscopy (EM) (Figure S7B), and no co-localization with the ER membranes could be observed (Figure S7C). Cytoplasmic nucleoporins were also found to be recruited to assembling SGs upon induction of cellular stress (Zhang et al., 2018). Consistently, our results demonstrated co-localization of the Nup RanBP2 with the markers of SGs, TIA-1 and G3BP1, in the control stress-induced cells (Figure S8A). However, TIA-1 and G3BP1 did not co-localize with the Nup aggregates in the FXR1-deficient cells exposed to stress, and both the SGs and Nup aggregates present in these cells localized to different cytoplasmic compartments. We conclude that the Nup aggregates in FXR1-deficient cells are distinct from ALs and SGs.

Given the established role of the FXR protein family in RNA-binding and the frequent role of RBPs in the formation and dynamics of membrane-less protein assemblies, we next analyzed whether Nup aggregates contain any RNAs or are linked to RNA-based processes. Hybridization with an RNA FISH probe against Poly-A revealed no difference in the percentage of nuclear mRNAs in the FXR1-deficient cells relative to control cells (Figure S8B, C), suggesting that the FXR1-Nup pathway is not implicated in mRNA export to the cytoplasm. Additionally, no cytoplasmic co-localization of the Nup aggregates and mRNAs could be observed in the FXR1-downregulated cells (Figure S8B, D). To test whether RNAs play a role in the maintenance or dynamics of the Nup aggregates in the cytoplasm, we treated permeabilized, FXR-downregulated cells with RNAse. This treatment failed to disrupt the aggregates or change their shape and distribution relative to control cells (Figure S9A, B), suggesting that RNAs are dispensable for their maintenance and dynamics. Collectively, we propose that these Nup aggregates represent previously unknown protein assemblies that form in the absence of FXR1 in human cells.

### FXR1 inhibits Nup aggregate formation by dynein-based microtubule-dependent transport

We next investigated the mechanism by which loss of FXR1 promotes the formation of Nup aggregates. We noticed that the aggregates were not scattered randomly in the cytoplasm but often formed a crescent-like shape around the microtubule-organizing centre (MTOC) suggesting a role for microtubules (Figure S10A). Indeed, we found that nocodazole-mediated microtubule depolymerization also induced Nup aggregate formation (Figure S10B, C). Notably, our earlier mass spectrometry analysis identified the cytoplasmic minus-end directed motor protein dynein heavy chain (HC) co-immunoprecipitating specifically with GFP-FXR1, along with the Nups (Table 1). We demonstrated an interaction of GFP-FXR1 with other components of the dynein complex, specifically dynein intermediate chain (IC) (which was visualized as a slower migrating band relative to the input lysate) and dynactin p150^Glued^ (Figure 4A). Downregulation of dynein HC by two independent siRNAs, which also depleted dynein IC as reported (Splinter et al., 2010) (Figure S10D), also led to the accumulation of the cytoplasmic Nup aggregates (Figure 4B, C and Figure S10E), highly reminiscent of FXR1 depletion. Analysis of several known dynein adaptor proteins in a co-immunoprecipitation assay with GFP-FXR1 revealed an interaction with BICD2, but not Mitosin or HOOK3 (Figure S10F). Downregulation of BICD2 by two independent siRNAs likewise led to accumulation of the cytoplasmic Nup aggregates (Figure S10G, H). Our results suggest that FXR1, working together with the microtubule motor dynein-BICD2 complex, are required to prevent cytoplasmic Nup aggregate formation.

**Figure 4.**
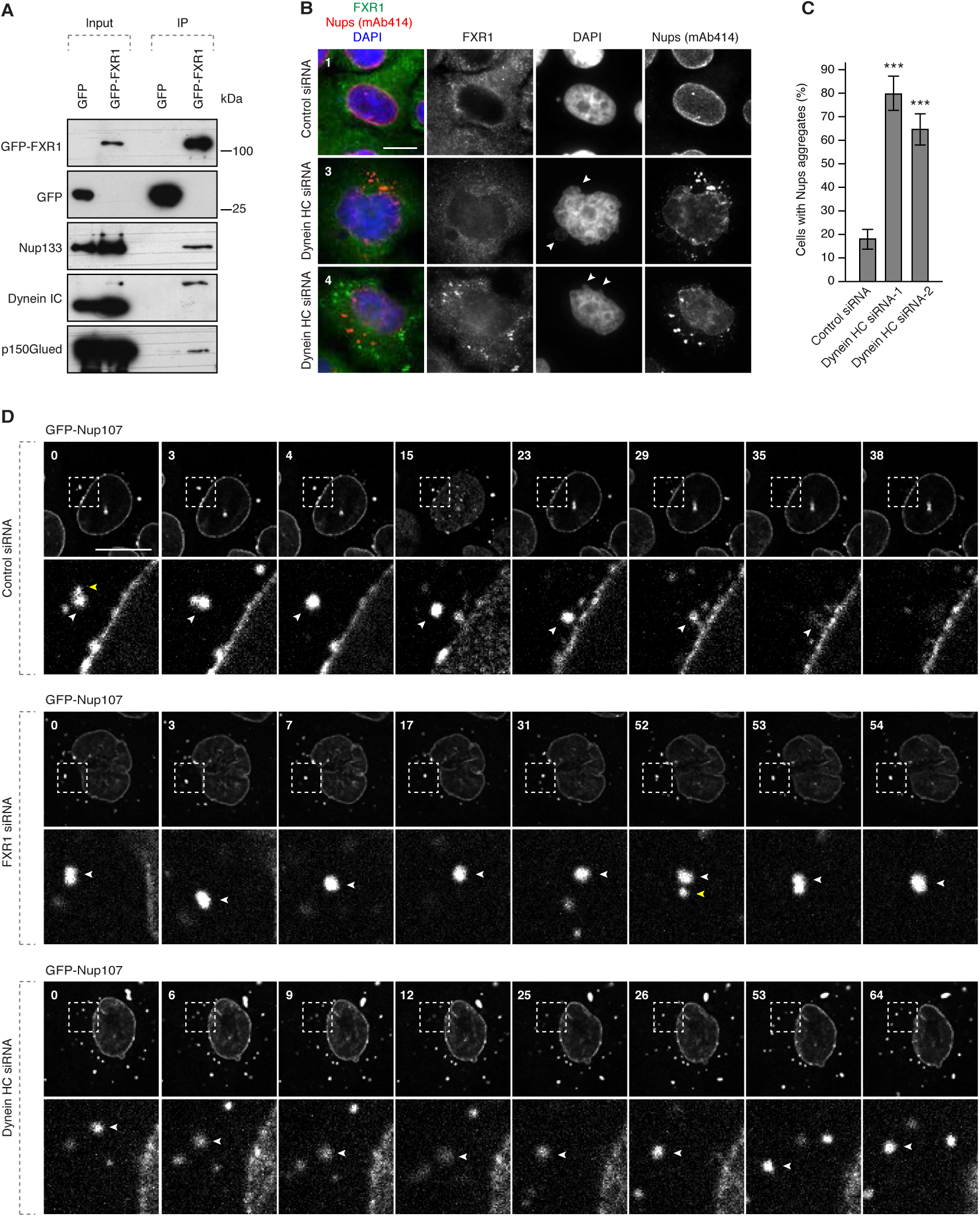
FXR1 regulates cytoplasmic Nups by dynein-based microtubule-dependent transport. **(A)** HeLa cells stably expressing GFP alone or GFP-FXR1 were immunoprecipitated using GFP-Trap beads (GFP-IP) and analysed by Western blot. (**B-C)** HeLa cells were treated with the indicated siRNAs, synchronized by double thymidine block, released for 12 hours and analysed by immunofluorescence microscopy. Images in (B) correspond to the numbered magnified framed regions indicated in the pictures shown in Figure S10E. The percentage of interphasic cells with aggregates (C) (n=900) was quantified. The corresponding Western blot analysis is shown in Figure S10D. Arrowheads indicate blebbed or herniated regions of nuclei. **(D)** HeLa cells stably expressing GFP-Nup107 were treated with the indicated siRNAs, synchronized by double thymidine block, released for 12 hours, treated with nocodazole to induce aggregate formation, washed-out and analysed by live video spinning disk confocal microscopy. The selected frames of the movies are depicted and time is shown in minutes. The magnified framed regions are depicted in the lower rows. White arrowheads point to individual GFP-Nup107-positive aggregates. Yellow arrowheads point to the aggregates undergoing fusion events. Bars are 5 µm.

Our earlier results in FXR1-deficient cells showed cytoplasmic Nup aggregate formation concomitant with a decrease in Nup localization at the NE. These results suggest that at least some of the Nups are no longer incorporated into the NE during interphase, and instead increase in concentrations in the cytoplasm where they ultimately form aggregates. Thus, we considered that FXR1 together with the microtubule motor dynein-BICD2 complex may mediate the transport of cytoplasmic Nups to the NE during interphase. To test this, we performed live video spinning disc microscopy of cell lines stably expressing GFP-Nup107, which is one of the Nups found in the aggregates. We then treated the cells with nocodazole to induce Nup aggregate formation in a reversible way. Following nocodazole wash-out, we observed the dynamics of the Nup aggregates in control, FXR1- and dynein-downregulated cells. GFP-Nup107-positive aggregates showed dynamic behavior and both fusion and splitting of the aggregates were observed under all conditions (Figure 4D). In control cells, nocodazole wash-out led to NE-directed transport and fusion of GFP-Nup107-aggregates with the NE. In contrast, downregulation of FXR1 or dynein in nocodazole washed-out cells led to retention of the GFP-Nup107-aggregates in the cytoplasm. Under these conditions, the GFP-NUP107-aggregates were still mobile but no NE-directed movement was observed, and they continued to fuse and often increased in size. These observations suggest that NE-directed transport by the FXR1-dynein complex can decrease local concentrations of cytoplasmic Nups thereby preventing their assembly into aggregates.

### Nup localization defects can be linked to Fragile X syndrome

Next, we analysed whether all members of the FXR protein family share analogous roles in the spatial control of Nup assembly. Our data show that in addition to FXR1, FXR2 and FMRP can localize at the NE in HeLa cells (Figure S11A) and in mouse myoblasts (Figure S11B). Interestingly, depletion of each of the three members of this protein family led to the aggregation of cytoplasmic Nups relative to control cells (Figure S11C, D). Simultaneous downregulation of all three FXR proteins did not further increase the penetrance of this phenotype (Figure S11C, D), suggesting that FXR proteins together form a protein complex in human cells consistent with our mass spectrometry results (Table 1). Since FMRP is silenced in Fragile X syndrome (Santoro et al., 2012), we asked if the Nup localization defects are observed in cellular models of this disease. We stimulated FXS patient-derived fibroblasts which lack the FMRP protein (Figure 5A) to undergo synchronous mitotic exit and nuclear reformation. FXS fibroblasts, but not control human fibroblasts, displayed accumulation of cytoplasmic Nup aggregates similar to those observed in HeLa cells (Figure 5B, C), and structurally abnormal nuclei (Figure 5D). To corroborate these findings, we used human induced pluripotent stem cells (iPSCs) derived from an FXS patient (FXS-iPSCs) and the isogenic rescue cells (C1_2-iPSCs), where reactivation of the *FMR1* locus is achieved by CRISPR-mediated excision of the expanded CGG-repeat from the 5’UTR of the *FMR1* gene (Xie et al., 2016). In the FXS-iPSCs, accumulation of large Nup133-positive cytoplasmic condensates was observed (Figure 5E), which were significantly reduced in the FMRP re-expressing cells, although they were still present (Figure 5E, F). Re-expressed FMRP in the reactivated cell line localized to both the NE and to the cytoplasmic perinuclear region, which often also contained Nup133 (Figure 5G). We speculate that these perinuclear FMRP-Nup133 signals could represent assembly intermediates before Nup133 is properly transferred and inserted into the NE. Thus, the absence of human FMRP in primary cell lines and in iPS cells leads to the accumulation of Nup aggregates as also seen in the cancer cell lines.

**Figure 5.**
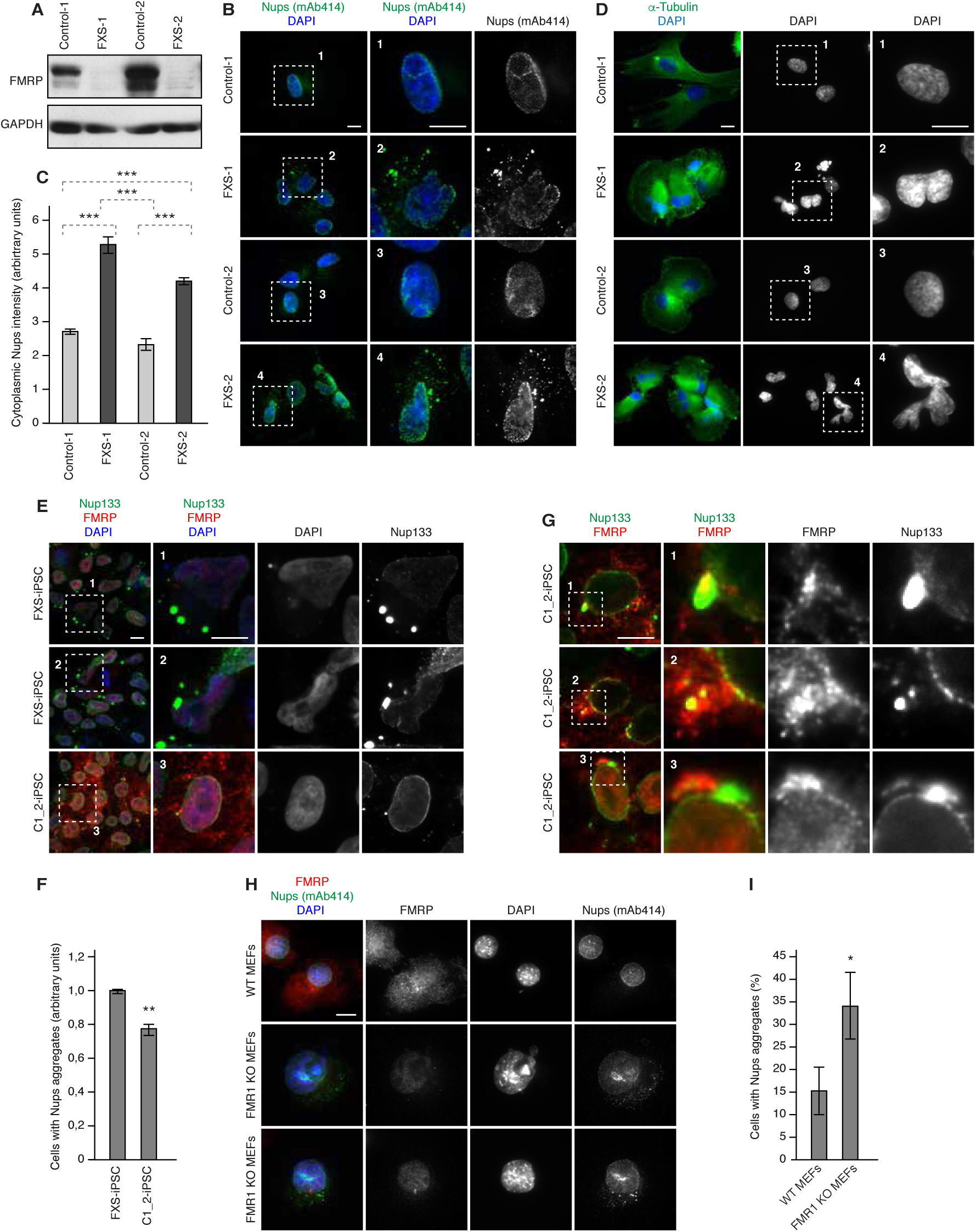
Nup localization defects can be linked to Fragile X syndrome. **(A-D)** Human FXS-patient derived fibroblasts (FXS-1, FXS-2) and control human fibroblasts were synchronized in early G1 by Monastrol release and analysed by Western blotting (A) and immunofluorescence microscopy (C-D). The cytoplasmic Nup intensity was measured in control and FXS fibroblasts (C) (n=283). Examples of Nup localization defects are shown in (B) and examples of nuclear architecture defects are shown in (D). **(E-G)** Human induced pluripotent stem cells (iPSCs) derived from a FXS patient (FXS-iPSC) and the isogenic rescue cells (C1_2-iPSC) were analysed by immunofluorescence microscopy. A ratio of cells with Nup aggregates was quantified in (n=5500) (F). Examples of co-localization events of re-expressed FMRP and Nup133 are shown in (G). **(H-I)** Mouse Embryonic Fibroblasts (MEFs) derived from the *FMR1* knock-out (KO) mice and wild type controls were synchronized in early G1 by Monastrol release and analysed by immunofluorescence microscopy. The percentage of cells with cytoplasmic nucleoporin aggregates (n=2400) was quantified in (I). Bars are 5 µm.

To confirm these findings in an animal model of FXS, we used mouse embryonic fibroblasts (MEFs) derived from the *FMR1* knock-out (KO) mice. *FMR1* KO MEFs also displayed accumulation of perinuclear Nup aggregates relative to wild type MEFs (Figure 5H, I). Taken together, our results demonstrate the presence of Nup assemblies in several cellular models of Fragile X syndrome. These defects may perturb cellular homeostasis and contribute to FXS pathology.

### The FXR protein family regulates nuclear morphology and G1 cell cycle progression

What could be the biological consequences of misregulation of the FXR-dynein pathway and how could Nup assembly defects perturb cellular homeostasis? FXR1 downregulation did not affect distribution of the nuclear import factor importin β1 (Figure S12A, B) and did not change the rates of nuclear import (Figure S12C, D) and export (Figure S12E, F) relative to control cells, whereas downregulation of the Nup ELYS clearly demonstrated import and export defects in the same experiments, as expected (Figure S12C-F). This is consistent with the earlier mRNA export results (Figure S8B, C) and indicates that, at least in the steady-state, nucleocytoplasmic transport is largely unaffected by formation of Nup aggregates in FXR1-deficient cells. These results are not surprising given relatively small reduction of the Nup signals at the NE upon FXR1 downregulation (Figure 2D-F) in interphase cells.

We noticed that the aggregate-containing cancer (Figures 1, 2, 4 and Figures S1-5, S7-10, 12) and primary cells (Figure 5) often displayed strong nuclear atypia. Indeed, downregulation of FXR1 led to defects in nuclear architecture including irregular, blebbed, and herniated nuclei (Figure 6A), which could be largely rescued by stable ectopic expression of the siRNA-resistant form of GFP-FXR1 (Figure 6B, C). Likewise, downregulation of dynein (Figure 6D) and all three FXR proteins (Figure 6E) led to nuclear morphology defects. Minor but detectable increases in nuclear size could also be measured in the FXR1-downregulated cells relative to control cells (Figure 6F). Live video microscopy revealed that the nuclear morphology defects in FXR1-deficient cells could first be detected approximately 30 min after the onset of chromosome decondensation (Figure 6G, H), which strongly correlated with the onset of nuclear growth and with the time when the first cytoplasmic Nup aggregates could be detected in the cytoplasm (Figure 3A, C). Consistent with the previous results (Figure 3A, C, Figure S6A, B), the timing of mitotic progression was not affected in the FXR1-deficient cells in this analysis (Figure 6I). Our data suggest that disruption of the FXR-dynein pathway specifically affects cytoplasmic Nup assembly and nuclear architecture during early G1. These observations prompted us to test whether the FXR pathway could also be important for G1 cell cycle progression. Downregulation of FXR1 in the asynchronously proliferating U2OS cell line stably expressing the FUCCI cell cycle sensor led to accumulation of cytoplasmic aggregates and reduction of nuclear Nup intensity, consistent with the results in the other cell lines (Figure 7 A-C). In addition, FXR1-downregulation led to an increase in the number of cells in G1- and a decrease in the number of cells in S/G2 phases (Figure 7A, C). These results were confirmed by analysis of a marker of the G0/G1 phase transition, phosphorylated retinoblastoma (Rb) protein (phospho-Rb), where downregulation of FXR1 led to accumulation of cells with strong nuclear phospho-Rb signal (Figure 7E, F). Together, these data indicate a delay in G1/S cell cycle progression in the absence of FXR1.

**Figure 6.**
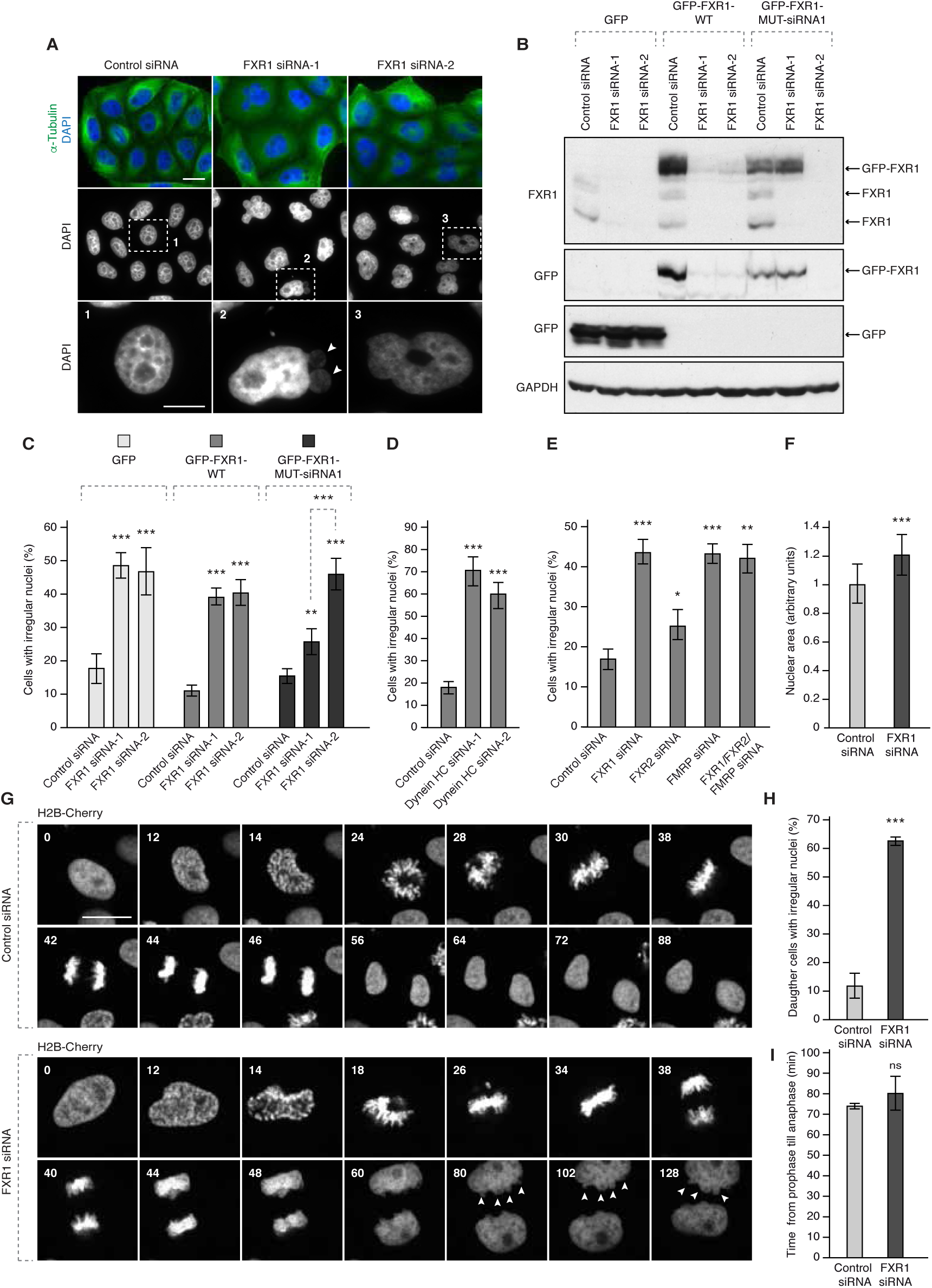
The FXR protein family control nuclear morphology. **(A)** HeLa cells were treated with the indicated siRNAs, synchronized by double thymidine block, released for 24 hours and analysed by immunofluorescence microscopy. The magnified framed regions are shown in the corresponding numbered panels. **(B-C)** HeLa cells stably expressing GFP, GFP-FXR1 wild type (WT) and GFP-FXR1 mutated in the sequence recognized by FXR1 siRNA-1 (GFP-FXR1-MUT-siRNA1) were treated with the indicated siRNAs, synchronized by double thymidine block, released for 24 hours and analysed by Western blotting (B) and immunofluorescence microscopy (C). The percentage of cells with irregular nuclei (n=1000) was quantified (C). The corresponding representative pictures are shown in Figure S1. **(D)** HeLa cells were treated with indicated siRNAs, synchronized by double thymidine block, released for 12 hours and analysed by immunofluorescence microscopy. The percentage of interphase cells with irregular nuclei (n=900) was quantified. **(E)** HeLa cells were treated with indicated siRNAs, synchronized by double thymidine block, released for 24 hours and analysed by immunofluorescence microscopy. The percentage of cells with irregular nuclei (n=1000) was quantified. **(F)** HeLa cells were treated and analysed as in (E). Nuclear area was quantified from 17 different biological replicates. **(G-I)** HeLa cells stably expressing the chromatin marker histone H2B labelled with mCherry were treated with indicated siRNAs, synchronized by double thymidine block, released and analysed by live video spinning disk confocal microscopy. The selected frames of the movies are depicted and time is shown in minutes (G). Arrowheads point to nuclear hernia and blebs appearing during nuclear expansion of FXR1-deficient cells. Percentage of daughter cells with irregular nuclei was quantified in (H) and time from prophase till anaphase was quantified in (I) (n=66). Bars are 5 µm.

**Figure 7.**
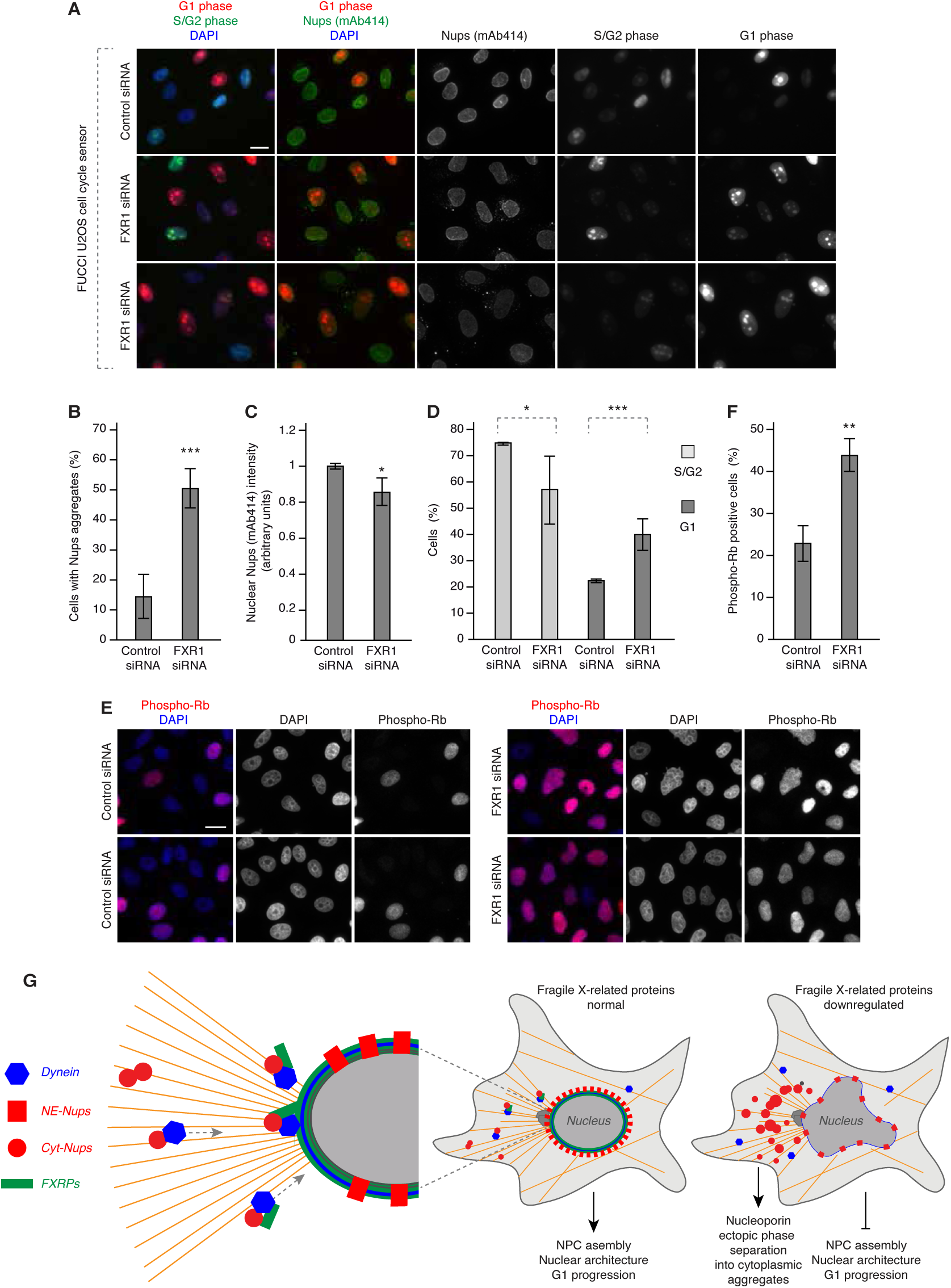
FXR1 regulates G1 cell cycle progression. **(A-D)** Asynchronously proliferating U2OS cells stably expressing the FUCCI cell cycle sensor were treated with indicated siRNAs and analysed by immunofluorescence microscopy. The percentage of cells with cytoplasmic nucleoporin aggregates (n=1600) was quantified in (B) and the nuclear FG-Nups intensity (n=1500) in (C). The percentage of cells in the G1-phase and S/G2 phase of the cell cycle (n=1600) was quantified in (D). **(E-F)** Asynchronously proliferating HeLa cells were treated with indicated siRNAs and analysed by immunofluorescence microscopy. The percentage of Phospho-Rb positive cells (n=2800) (corresponding to G1-phase of the cell cycle) was quantified in (F). **(G)** A model for how FXR family proteins regulate cytoplasmic Nups. FXR proteins (green) interact with the excess of the cytoplasmic Nups (red circles) and dynein (blue) and facilitate their localization to the NE during early G1. This chaperonin function of FXR proteins inhibits formation of aberrant cytoplasmic Nup condensates, the CNGs, contributing to the equilibrium of NE-NPCs (red rectangles) and to the maintenance of nuclear shape and G1 cell cycle progression. Silencing of FXR proteins, for instance in FXS patients, leads to the accumulation of CNGs, nuclear atypia and a delay in G1 progression, which may contribute to the pathology of FXS.

## Discussion

Collectively, our data suggest a model where FXR proteins and dynein regulate the localization of a cytoplasmic pool of Nups thereby facilitating their NE assembly during early G1 (Figure 7G) and decreasing cytoplasmic Nup concentrations. Absence of FXR proteins or dynein-mediated transport leads to the formation of previously uncharacterized ectopic Nup assemblies. We speculate that the FXR-dynein pathway regulates the pool of nucleoporins remaining in the cytoplasm after post-mitotic NPC assembly, which is important for G1-specific functions in nuclear shape control and cell cycle progression. Defects in this pathway, as seen in cellular models of Fragile X syndrome, may therefore compromise cellular fitness and contribute to the pathology of this incurable human disease.

Our analysis demonstrates that FXR proteins can act as molecular chaperones that facilitate dispersal of Nups and reversal of cytoplasmic Nup assemblies, which we propose to name Cytoplasmic Nucleoporin Granules (CNGs) (Figure 7G). Consistent with this hypothesis, FXR1 in cancer cells (Figure 1E) and re-expressed FMRP in IP cells (Figure 5G) could both co-localize in foci with a pool of cytoplasmic Nups often found in the proximity of the NE. FXR1could also interact with Nups and rescue experiments with the siRNA resistant form of FXR1 could reverse the formation of CNGs in cancer cells (Figure 2B and Figure S1). Finally, the CNGs fusion events observed in the nocodazole wash-out experiments, resulted in increase of the CNGs’ size in the absence of FXR1 or dynein relative to control cells (Figure 4D).

The CNGs formed in the absence of FXR proteins likely represent distinct structures from ALs based on our EM analysis (Figure S7B) and lack of direct contacts with the ER membranes (Figure S7C). However, it cannot be excluded that some membranes or lipid species are part of the CNGs due to the presence of the integral membrane proteins POM121 and Gp210, both components of the NPC at the NE, and the scaffold NPC component Nup133 in these structures. The CNGs formed in the absence of FXR proteins are also distinct from stress granules (SGs) (Figure S8A) previously shown to recruit nucleoporins (Zhang et al., 2018) and are not directly linked to RNA-based processes (Figure S8B-D, S9). We were unable to detect any changes in Nup protein levels (Figure S6A-C) or the levels and stability of several Nup mRNAs (Figure S6D, E) excluding the possibility that FXR proteins are involved in their translational regulation, which represents one of the well-studied roles of the FXR protein family (Ascano et al., 2012; Darnell et al., 2009). Downregulation of FXR proteins did not inhibit the recruitment of other nuclear membrane proteins to the NE (Figure S2A-G), suggesting that cytoplasmic Nup condensates are not the result of defects in assembly of the nuclear lamina. Our data suggest a model where FXR proteins interact with the cytoplasmic pool of the scaffold NPC components, Nup85 and Nup133 and act to decrease their local concentration by transfering them towards the NE. Due to the cohesive ability of many Nups and the fact that the scaffold components can directly bind to the FG-Nups (Onischenko et al., 2017), it is therefore reasonable to predict that in the absence of the FXR proteins, cytoplasmic levels of Nup85 and Nup133 rise, and could result in the formation of cytoplasmic condensates containing many different NPC sub-complexes, which is consistent with our observations (Figures 1, 2, 4 and Figures S3-5, S7). It remains to be investigated if the absence of ELYS and Nup153 in the CNGs is linked to their well-established roles in the post-mitotic (ELYS) (Doucet et al., 2010) and interphase (Nup153) (Vollmer et al., 2015) nuclear pore assembly pathways.

What is the molecular engine for the FXR-chaperonin function? Interestingly, FMRP was demonstrated to form a complex with the dynein motor (Bianco et al., 2010; Ling et al., 2004) and with the dynein adaptor protein BiCD (Bianco et al., 2010) in neuronal cells. Our data are consistent with these findings and show the interaction of dynein and BICD2 with the FMRP paralog protein FXR1 (Figure 4A, Figure S10F) in cultured human cancer cells. Molecular interactions of nucleoporins and dynein-BICD2 complexes were also reported during mitotic entry (Bolhy et al., 2011; Splinter et al., 2010). Our model proposes that FXR proteins provide the molecular links between cytoplasmic Nups and the dynein-BICD2 complex during the G1 phase of the cell cycle and, after successful completion of post-mitotic nuclear pore assembly, allow for the transfer of excessive scaffold Nups towards the NE. It is interesting that FXR proteins localize to the NE (Figure 1D-F, Figure S11A, B), the predicted final destination of the cytoplasmic Nups. Our live video experiments are in line with the transport hypothesis and suggest that nocodazole-induced CNGs can indeed be transferred towards intact NE (Figure 4D) in an FXR1- and dynein-dependent manner. We speculate that formation of FXR1-dynein-Nup complexes and their transport would disperse cytoplasmic nucleoporins, thereby inhibiting formation of Nup-containing cytoplasmic condensates.

A similar function has been ascribed to the nuclear import receptors, which transfer proteins and RNAs to the nucleus across the NPCs. They were able to prevent aberrant phase separation of cytoplasmic membrane-less organelles present in several neurological diseases (Guo et al., 2019). Thus, nuclear import receptors can also play important chaperone-like functions by inhibiting aggregation of cargo proteins. For instance, protein fused in sarcoma (FUS) is mutated in amyotrophic lateral sclerosis (ALS) within its nuclear localization signal (NLS), which subsequently reduces its binding to the nuclear import receptor Transportin leading to cytoplasmic FUS accumulation favoring phase separation (Dormann et al., 2010). Our work also demonstrates the importance of the regulation of localized protein demixing by facilitating Nup assembly at the NE and preventing their assembly in the wrong cellular compartment (cytoplasm). In the future, it would be interesting to study if the NLS signals present in the FXR proteins are important for their chaperone functions on Nups. It also remains to be investigated if, in contrast to stress granules components (Figure S8A) (Zhang et al., 2018), CNGs can sequester any other cohesive proteins known to form membrane-less assemblies. For example Nups can be sequestered in various pathological fibrillary amyloids, which were implicated in neurodegenerative diseases (Hutten and Dormann, 2019; Li and Lagier-Tourenne, 2018) and in the CyPNs (cytoplasmic accumulations of PML and nucleoporins) (Jul-Larsen et al., 2009), for which the cellular role remains unknown.

Our data suggest that the FXR-dynein pathway is important for the maintenance of nuclear shape during early G1. Downregulation of all members of the FXR family and dynein led to a strong defect in nuclear shape in human cancer cells but nuclear atypia has been also observed in human primary fibroblasts derived from FXS patients, as well as in the FMRP-deficient IPS cells and MEFs. These results are in line with the established structural functions of nucleoporins ensuring correct nuclear shape (Grossman et al., 2012). Indeed, changes in nuclear shape in cells deficient for individual Nups have been documented in various organisms (Hetzer and Wente, 2009; Mitchell et al., 2010; Onischenko et al., 2017; Ungricht et al., 2015). The first changes in nuclear morphology are observed during early G1, which correlates with the appearance of CNGs in the FXR1-deficient cells and may consequently delay the progression of the cell cycle throught G1. Interestingly, the components of the Y-complex were previously implicated in G1/S progression by regulating export of specific mRNAs of key cell cycle genes (Chakraborty et al., 2008). Our data do not show global changes in the rates of export and import (Figure S12) under normal conditions, which is expected given the relatively small reduction of NE-associated Nups observed in FXR1-downregulated cells (Figure 2D-F). Nevertheless, it cannot be excluded that under stress conditions or in the fast dividing cells of a developing embryo, this small decrease of NE-Nups would impact nucleocytoplasmic exchange. It is also possible that the transport of specific or more demanding cargos through NPCs is regulated by FXR-dynein pathway. For instance in yeast, modulation of NPCs in the daughter cells drives their cell cycle entry by specifically regulating nuclear export of Whi5 (Rb protein in humans) (Kumar et al., 2018). Our data in the FXR1-deficient cells show accumulation of cells with phosphorylated nuclear Rb (P-Rb) (Figure 7E, F), which supports the speculation that FXR1-dependent G1 regulation could be due to specific effects on P-Rb export. An alternative and equally plausible explanation is that CNGs exert cytotoxic effects by sequestering yet unknown factors important for nuclear shape and cell cycle progression. While future studies are needed to understand the precise mechanism underlying G1 control by the FXR-dynein axis, and if and how it is linked to the regulation of cytoplasmic Nups, defects in this pathway are predicted to significantly perturb cellular homeostasis and may contribute to the pathology of Fragile X syndrome, consistent with our observations in the cellular models of FXS (Figure 5). Collectively, our data demonstrate an unexpected role of FXR proteins and dynein in the spatial regulation of Nups, and provide an example of a mechanism that regulates localized protein condensate assembly.

## Acknowledgements

We thank patients and their families for their contribution. Y. Barral, S. Oliferenko, G. Sumara, O. Sumara, M. Mendoza, N. Djouder, K. Krupina, O. Bielska, Z. Zhang, Life Science Editors and the members of the Sumara group for helpful discussions on the manuscript. We are grateful to Valérie Doye for generous help with reagents. We thank Ulrike Kutay, Frauke Melchior, Hélène Puccio, Jin Peng, Stephen Warren, Romeo Ricci, Jan M. van Deursen and Nicolas Charlet-Berguerand for help with reagents. We are grateful to Jean-Marc Egly and Michel Labouesse for their mentorship and support. We thank the Imaging Center of the IGBMC (ICI) for help on electron and confocal microscopy and to the IGBMC core facilities for their support on this research. Alexia Loynton-Ferrand from the Biozentrum, University of Basel for support on superresolution imaging. A.A.A. is supported by a Labex international PhD fellowship from IGBMC and a fellowship from the “Ligue Nationale Contre le Cancer”. S.S. was supported by a postdoctoral fellowship from the University of Strasbourg Institute of Advanced Studies (USIAS). K.J. is supported by a fellowship from Gouvernement français et L’Institut français de Prague, a LabEx international PhD fellowship from IGBMC and a fellowship from the “Ligue Nationale Contre le Cancer”. A.B. received PhD fellowships from the “Ministère de l’Enseignement Supérieur et de la Recherche” and the “Ligue Nationale contre le Cancer”. This study was supported by the grant ANR-10-LABX-0030-INRT, a French State fund managed by the Agence Nationale de la Recherche under the frame program Investissements d’Avenir ANR-10-IDEX-0002-02. Research in I.S. laboratory was supported by IGBMC, CNRS, Fondation ARC pour la recherche sur le cancer, Institut National du Cancer (INCa), Agence Nationale de la Recherche (ANR), Ligue Nationale contre le Cancer, USIAS and Sanofi iAward Europe.

## Author contributions

A.A.A. and S.S. designed and performed experiments and helped writing the manuscript. K.J., C.K., A.B, L.P. I.J.B, P.R. and L.G. performed experiments. H.M. helped performing experiments. S.J. and C.B. provided human patient samples. J.L.M. and Y.S. helped designing the experiments. I.S. supervised the project, designed experiments and wrote the manuscript with input from all authors.

## Additional information

**Supplementary information** is available for this paper at https://

**Correspondence and requests for materials** should be addressed to I.S.

## STAR Methods

### Cell lines and cell cycle synchronizations

HeLa Kyoto and derived stable cell lines (GFP, GFP-FXR1, 3xGFP-NUP85, GFP-NUP107) were cultured in Dulbecco’s modified Eagle Medium (DMEM) (4,5 g/L glucose, with GLUTAMAX-I) supplemented with 10% FCS, 1% Penicillin and 1% Streptomycin. Cells were synchronized by two-times addition of Thymidine at 2 mM for 16h. Cells were washed out after each Thymidine addition three times with warm medium to allow for synchronous progression through cell cycle. Cells were analysed at desired time points after the release from the second thymidine block. Alternatively, cells were synchronized in early G1 by inducing an artificial mitotic exit. First, cells were treated with Taxol (paclitaxel) for 16h at 1 µM and then subsequently released from the mitotic block by addition of Hesperadin at 100 µM for 2h. Human primary fibroblasts were cultured in DMEM (4,5 g/L glucose) supplemented with 10% FCS and Gentamicin 40 µl/ml. Three independent mouse embryonic fibroblasts (MEFs) lines from control and three MEFs from *FMR1* knockout mice were cultured in DMEM (4,5 g/L glucose) supplemented with 10% FCS, 1% Penicillin and 1% Streptomycin. Fibroblasts and MEFs were synchronized with 100 µM Monastrol (Sigma, M8515) for 16h, washed three times with warm medium and released into fresh medium for 2h. HEK293T cells were cultured asynchronously in Dulbecco’s modified Eagle Medium (DMEM) (1 g/L glucose) supplemented with 10% FCS and 1x Penicillin, Streptomycin. U2OS cells stably expressing FUCCI cell cycle sensor (Macurek et al., 2013) were cultured asynchronously in DMEM (4,5 g/L glucose, with GLUTAMAX-I) supplemented with 10% FCS, 1% Penicillin and 1% Streptomycin. Mouse myoblasts (C2C12) were cultured asynchronously in DMEM (1 g/L glucose) supplemented with 20% FCS and gentamicin. Human induced pluripotent stem cells (iPSCs) derived from a FXS patient (FXS-iPSCs) and the isogenic rescue cells (C1_2-iPSCs) were grown asynchronously as indicated by (Xie et al., 2016).

### Immunofluorescence microscopy and sample preparation

Cells were plated on 9-15 mm glass coverslips (Menzel-Glaser) in 12- or 24-well tissue culture plates. At the end of the experiments cells were washed twice with PBS and fixed for 10 minutes with 1% paraformaldehyde (PFA) in PBS at room temperature (RT). The coverslips were rinsed two times with PBS and permeabilized with 0.1% Triton X-100 and 0.02% SDS in PBS for 5 minutes at room temperature, washed two times with PBS and blocked by blocking buffer 3% BSA/PBS-T (0.01% Triton X-100). Coverslips were subsequently incubated with primary antibodies in blocking buffer for 1 hour at room temperature, rinsed three times with blocking buffer and incubated with secondary antibodies in blocking buffer for 30 min at room temperature in the dark. After incubation, coverslips were rinsed three times with blocking buffer and mounted on glass slides using Mowiol (Calbiochem) or Prolong Gold reagent (Invitrogen) with 0.75 µg/µl DAPI and imaged with a 100X, 63X or 40X objectives using Zeiss epifluorescence microscope or confocal microscope Leica Spinning Disk Andor/Yokogawa. For experiments with human fibroblasts and MEFs, cells were fixed with 4% PFA for 17 minutes, washed 3 times in PBS and permeabilized with 0.5% NP-40 in PBS for 5 minutes. Cells were washed 3 times with PBS-T and blocked in 3% BSA/PBS-T for 1 hour. Cells were incubated with primary antibodies for 90 minutes in blocking buffer, washed 3 times in PBS-T and incubated with secondary antibodies for 1 hour. Cells were washed 3 times in PBS-T and mounted as previously described. For digitonin permeabilization experiments, cells with treated as indicated but permeabilized with 0,003% digitonin in PBS for 5 minutes and subsequent steps were performed without detergent. To induce formation of the cytoplasmic nucleoporin aggregates by microtubule depolymerization, cells were incubated with 10 μM Nocodazole or vehicle (DMSO) in culture media for 90 minutes at 37 °C. Subsequently, immunofluorescence protocol was performed as previously described. To observe the behavior of cytoplasmic nucleoporin aggregates upon microtubule repolymerization in live-video microscopy, nocodazole was washed 5 times with warm media during image acquisition.

### Poly A RNA Fluorescent In Situ Hybridization (FISH)

HeLa K GFP-Nup107 cells were plated on 9-15 mm glass coverslips (Menzel-Glaser) in 12- or 24-well tissue culture plates. At the end of the experiments cells were washed once with PBS and fixed for 10 minutes with 4% PFA in PBS at RT. Subsequently, cells were incubated with 100% cold methanol at −20 °C for 10 minutes. The coverslips were incubated with 70% ethanol at 4 °C overnight. Cells were incubated in 1M Tris-HCl pH 8 for 5 minutes at RT before proceeding to hybridization for 3 hours at 37 °C in hybridization buffer (2x SSC (Saline Sodium Citrate buffer), 1 mg/mL yeast tRNA, 0,005% BSA, 10% dextran sulfate, 25% formamide, 1 ng/uL oligo(dT_30_) fluorescent probes fused to Atto-565 or Atto-488 (Sigma)) protected from light. After hybridization, cells were washed once with 4x SSC, and 2 times with 2x SSC. Coverslips were mounted and imaged as indicated previously. For RNase treatment experiments, cells were washed 3 times with PBS and permeabilized with our without RNase A/T1 (0,2 mg/mL, 500 u/mL) in 0,003% digitonin PBS for 5 minutes, before performing the FISH protocol as described previously. All buffers were DEPC treated before use.

### Microscopy and image analysis

For live-cell microscopy, HeLa cells stably expressing indicated proteins tagged with GFP or mCherry, were grown on LabTek II Chambered Slides (Thermo Scientific) or µ-Slide VI 0.4 (IBIDI) or 35/10 mm glass bottom dishes (Greiner Bio-One, 627871). Before filming, cells were treated with SiR-DNA or Sir-Tubulin probes following manufacturer’s instructions when indicated. Live-cell microscopy was carried out using 40X or 63X objective of confocal microscope Leica/Andor/Yokogawa Spinning Disk or Leica CSU-W1 spinning disc.

For protein import assay, HeLa cells were treated with the indicated siRNAs for 72 hours and transfected with the reporter plasmid XRGG-GFP (kindly provided by Jan M. van Deursen) (Hamada et al., 2011; Love et al., 1998) 30 hours before filming. Cells were synchronized in early G1 phase by 100 μM Monastrol arrest for 16 hours and released for 4 hours. 1,25 μM dexamethasone-induced nuclear import of XRGG-GFP was recorded by live video spinning disk confocal microscopy for 25 minutes (1 acquisition every 30 seconds).

For protein export assay, HeLa cells were treated as described above and 0,25 μM dexamethasone was added for 3 hours to induce XRGG-GFP nuclear import. Following wash-out, the nuclear export of XRGG-GFP was recorded by live video spinning disk confocal microscopy for 2 hours (1 acquisition every 10 minutes).

For nucleoporin dynamics assays, HeLa cells stably expressing GFP-Nup107 were treated with the indicated siRNAs for 72 hours and synchronized in early G1 phase by 100 μM Monastrol arrest for 16 hours and released for 2 hours. Microtubule depolymerization and nucleoporin aggregation was induced by 10 μM nocodazole addition for 1,5 hours. Following wash-out, nucleoporin dynamics were recorded by live video spinning disk confocal microscopy for 90 minutes (1 acquisition every minute).

Image analysis was performed using ImageJ, CellProfiler or Metamorph software. Super-resolution microscopy was performed using API OMX “Blaze” with GE DeltaVision OMX stand and analysed with DeltaVision OMX softWoRx. Cells were grown on #1.5 High Precision Coverslips, fixed, permeabilized and stained according to the protocol for the fluorescent microscopy (see above). Coverslips were mounted onto the microscope slides with Vectashield H1000 mounting medium (soft setting) and sealed with a nail polish.

### Statistical analysis

Where indicated, analysis of the difference between two groups was performed with unpaired two-tailed t test (with p < 0.05). Error bars represent Standard Deviation (SD) except for live video experiments where bars represent Standart Error of the Mean (SEM). At least three independent biological replicates were performed for each experiment.

### Plasmid and siRNA transfections

Lipofectamine 2000 (Invitrogen) was used to deliver XRGG plasmid (kindly provided by Jan M. van Deursen) (Hamada et al., 2011; Love et al., 1998) according to the manufacturer’s instructions. Oligofectamine (Invitrogen) was used to deliver siRNAs for gene knockdown according to the manufacturer’s instructions at a final concentration of 40-100 nM siRNA. The following siRNA oligonucleotides were used: for non-silencing controls siGENOME Non-targeting siRNA Pool-1 and siGENOME Non-targeting individual siRNA-2 5′-UAAGGCUAUGAAGAGAUAC-3′ (Dharmacon), for FXR downregulation siRNA (Dharmacon) were used: FXR1 siRNA-1 5’-AAACGGAAUCUGAGCGUAA-3’; FXR1 siRNA-2 5’-CCAUACAGCUUACUUGAUA-3’; FXR2 5’-CGACAAGGCUGGAUAUAGC-3’; FMRP 5’-AAAGCUAUGUGACUGAUGA-3’; Dynein HC siRNA-1 5’-CGUACUCCCGUGAUUGAUG-3’; Dynein HC siRNA-2 5’-GGAUCAAACAUGACGGAAU-3’; BICD2 siRNA-1 5’-GGA GCU GUC ACA CUA CAU G-3’; BICD2 siRNA-2 5’-GGU GGA CUA UGA GGC UAU C-3’; ELYS siRNA 5’-AUU AUC UAC AUA AUU GCU CUU TT-3’. For endoplasmic reticulum observation experiments, cells were incubated with CellLight ER-RFP BacMam 2.0 (ThermoFischer, C10591) for 24h before following the manufacturer’s instructions.

### Quantitative Real-Time PCR

RNA of cultured cells was isolated using TRIzol reagent (Sigma) according to manufacturer’s instructions. Reverse transcription was performed with random hexamer or oligodT primers using the SuperScript III First Strand cDNA Synthesis kit (Invitrogen). SYBR Green (Roche Diagnostics) based Real-time PCR was carried out on the LightCycler 480 (Roche Diagnostics) using gene specific primer pairs:

NUP85: 5’-GACTGAACAAGTTCGCAGCA-3’ (forward), 5’-TCAGTCGGTCACTGAGCATC-3’ (reverse); GAPDH 5’-ACCCAGAAGACTGTGGATGG-3’ (forward), 5’-TTCTAGACGGCAGGTCAGGT-3’ (reverse); PO 5’-GTGATGTGCAGCTGATCAAGACT-3’ (forward) 5’-GATGACCAGCCCAAAGGAGA-3’ (reverse)

### Primers and molecular cloning

Cloning was performed using New England Biolabs (NEB) restriction enzymes, Taq polymerase (NEB) or Fusion High-Fidelity DNA polymerase (Thermo Scientific) according to the manufacturers’ instructions. To clone GFP-FXR1, FXR1 was amplified from HeLa total cDNAs with primers 5’-ttattaCTCGAGCCATGGCGGAGCTGACGGTGGAGG-3’ (forward) and 5’-tattatGAATTCTTATGAAACACCATTCAGGACTGC-3’ (reverse). FXR1 cDNA was cloned into the pEGFPC1 vector using EcoRI/XhoI restriction enzymes. To obtain the plasmid used in rescue experiment expressing GFP-FXR1-MUT-siRNA1, primers 5’-CTTTCCGTTCGCTCTCTGTCTCAGAGGGGTTAGACAGCTCAGAATTTG-3’ (forward) and 5’-TGAGACAGAGAGCGAACGGAAAGACGAGCTGAGTGATTGGTCATTGGC-3’ (reverse) to mutate pEGFPC1-FXR1 vector in the region targeted by the FXR1 siRNA1. The pEGFPC1-NUP85 was kindly provided by Valérie Doye. For live video microscopy experiements, cDNAs were cloned into a pVITRO-blasticidin vector (Invivogen). mCherry-H2B was amplified with primers 5’-AATAATGCTAGCATGCCAGAGCCAGCGAAGTCTGC-3’ (forward) and 5’-TAATAATCTAGATTACTTGTACAGCTCGTCCATGC-3’ (reverse). cDNA was cloned into the pVITRO in the multi cloning site 1 using NheI/XbaI restriction enzymes. GFP-NUP85 was amplified with primers 5’-aataatGCTAGCGCCATCATGGTGAGCAAGGGCGAGGAGCTG-3’ (forward) and 5’-tattatGCTAGCTCAGGAACCTTCCAGTGAGCCTTCTC-3’ (reverse). cDNA was cloned into the pVITRO in the multi cloning site 2 using the NheI restriction enzyme. DNA purifications were performed using commercial kits from Macherey-Nagel according to the manufacturer’s instructions.

### Generation or acquisition of stable cell lines

Hela cells were transfected with pVITRO or pEGFPC1-derived constructs using Lipofectamine 2000 according to manufacturer’s instructions. Transfected cells were selected for 2-3 weeks in medium supplemented by antibiotics, either G418 (400 µg/ml) or blasticidin (5 µg/ml). Transgene-expressing clones were then isolated by FACS (FACS ARIA, BD Biosciences). Expression was validated by Western blotting and immunofluorescence analysis. GFP-NUP107 stable cell line was purchased from CLS cell bank. U2OS cells stably expressing the FUCCI cell cycle sensor were kindly provided by Libor Macurek.

### Western blotting and immunoprecipitations

HeLa cell extracts were prepared using lysis buffer (50 mM Tris HCl pH 7.5, 150 mM NaCl, 1% Triton X-100, 1 mM EDTA 0.25% Sodium Deoxycholate, 1mM PMSF, protease inhibitor coctail). For GFP immunoprecipitation, GFP-fused proteins were immunoprecipitated using GFP-Trap A agarose beads (Chromotek). Beads were incubated with cell extracts for 1h30 at 4 °C under constant rotation. Before elution, beads were washed 3 times with washing buffer (10 mM Tris HCl pH 7.5, 150 mM NaCl, 0.5 mM EDTA, 0.1% Tween-20, protease inhibitor cocktail), boiled in Laemmli SDS sample buffer and subjected to SDS-PAGE.

For endogenous immunoprecipitation experiments, protein G sepharose 4 Fast Flow beads (GE Healthcare Life Sciences) were washed three times for 1 minute in washing buffer (Tris HCl 1M pH 7,5, NaCl 150 mM, EDTA 1 mM, protease inhibitor cocktail). FXR1 protein from HEK293T cell extracts were incubated with the beads and FXR1 antibody or rabbit IgG (1,5 uL per mg of protein) for 3 hours at 4 °C under constant rotation. Before elution, beads were washed 5 times during 2 hours with washing buffer 0.1% Tween-20, boiled in Laemmli SDS sample buffer and subjected to SDS-PAGE. Proteins were subsequently transferred from the gel to a PVDF membrane (Millipore) for immunoblotting. Membranes were blocked in 5% non-fat milk powder resuspended in TBS supplemented with 0.1% Tween 20 (TBS-T) for from 1 h to overnight, followed by incubation with antibodies. Membranes were developed with Luminata Forte (Millipore) or SuperSignal West Pico chemiluminescent substrate (Thermo Scientific).

### Electron microscopy

HeLa cells stably expressing 3xGFP-Nup85 were grown on carbon-coated sapphire disks and synchronized by double thymidine block and 12 hour release. After synchronization, cells were high pressure frozen (HPM010, AbraFluid) and freeze substituted with 0.1% Uranyl acetate in acetone for 15h. The temperature was then raised to −45°C at 5°C/h and cells were further incubated for 5h. After rinsing in acetone, the samples were finally embedded in Lowicryl HM20 (Polysciences Inc.). Thick sections (300nm) were cut from the UV-polymerized resin block and picked up on carbon coated mesh grids. After post-staining, 2D montages and tilt series of the areas of interest were acquired using a FEI TECNAI F30 TEM. Tomograms were reconstructed using the software package IMOD (Kremer et al., 1996).

### Antibodies

The following antibodies were used:

mouse monoclonal α-tubulin (Sigma T5169, immunofluorescence microscopy 1:4000, Western blotting 1:20000), rabbit α-GAPDH (Sigma G9545, Western blotting 1:20000), mouse α-GAPDH (Genetex, gtx627408, 1:20000), mouse monoclonal α-FXR1+2 (clone 2B12 from IGBMC, immunofluorescence microscopy 1:500, Western blotting 1:1000), rabbit α-FXR1 (Sigma HPA018246, immunofluorescence microscopy 1:800, Western blotting 1:1000), mouse α-FXR1 (Millipore 03-176, immunofluorescence microscopy 1:800, Western blotting 1:1000), mouse α-FMRP (clone 1C3 from IGBMC, immunofluorescence microscopy 1:250, Western blotting 1:1000), rabbit α-FMRP (Abcam ab17722, immunofluorescence microscopy 1:250, Western blotting 1:1000), mouse α-Nup133 (Santa cruz sc-37673, Western blotting 1:1000), rat α-Nup133 (kindly provided by Valérie Doye, immunofluorescence microscopy 1:250), mouse α-FG-Nups (Abcam mAb414, ab24609, 1:4000), rabbit α-Nup153 (Abcam ab84872, immunofluorescence microscopy 1:500, Western blotting 1:1000), rabbit α-Nup85 (Bethyl, A303-977A, immunofluorescence microscopy 1:100), mouse α-Pericentrin-1 (D-4) (NUP85) (Santa Cruz, sc-376111, Western blotting 1:1000), mouse α-PLK1 (Santa cruz sc-17783, Western blotting 1:1000), rabbit polyclonal GFP (Abcam ab290, 1:20000), mouse monoclonal Cyclin B1 (Santa Cruz sc-245, clone GSN1, 1:2000), rabbit α-cyclin A (Santa Cruz sc-751, 1:1000), rabbit α-laminA (Sigma L1293 1:500), rabbit α-lamin B1 (Abcam, ab16048, 1:500), mouse α-Lap2 (BD biosciences, 611000, 1:500,) rabbit α-ELYS (Bethyl, A300-166A, 1:250), rabbit α-POM121 (Genetex, GTX102128, 1:200), mouse α-RanGAP1 (Cell Signalling 2365, 1:250), rabbit α-emerin (Abcam, ab40688, 1:1000), rabbit α-LBR (Abcam, ab35535, 1:500), mouse α-Dynein IC (Merck, MAB1618, 1:500), mouse α-p150^glued^ (BD biosciences, 610473, 1:1000), goat α-RanBP2 (kindly provided by Frauke Melchior, 1:2000), rabbit α-Nup98 (Cell Signalling, 2598S, 1:100), rabbit α-BICD2 (Sigma HPA023013, immunofluorescence microscopy 1:250, Western blotting 1:1000), rabbit α-G3BP1 (Genetex GTX112191, immunofluorescence microscopy 1:500), mouse α-TIA-1 (G-3) (Santa Cruz, sc-166247, immunofluorescence microscopy 1:500), rabbit α-phospho-Rb (Cell Signaling 8516, immunofluorescence microscopy 1:1600), rabbit α-karyopherin β1 (H-300) (Importin β1) (Santa Cruz sc-11367, immunofluorescence microscopy 1:200). Antibodies against Nup88, Nup210, RAE1 were kindly provided by Ulrike Kutay.

## Data availability

All data supporting the findings of this study are available from the corresponding author upon reasonable request.

---

**Supplemental Figure 1:**
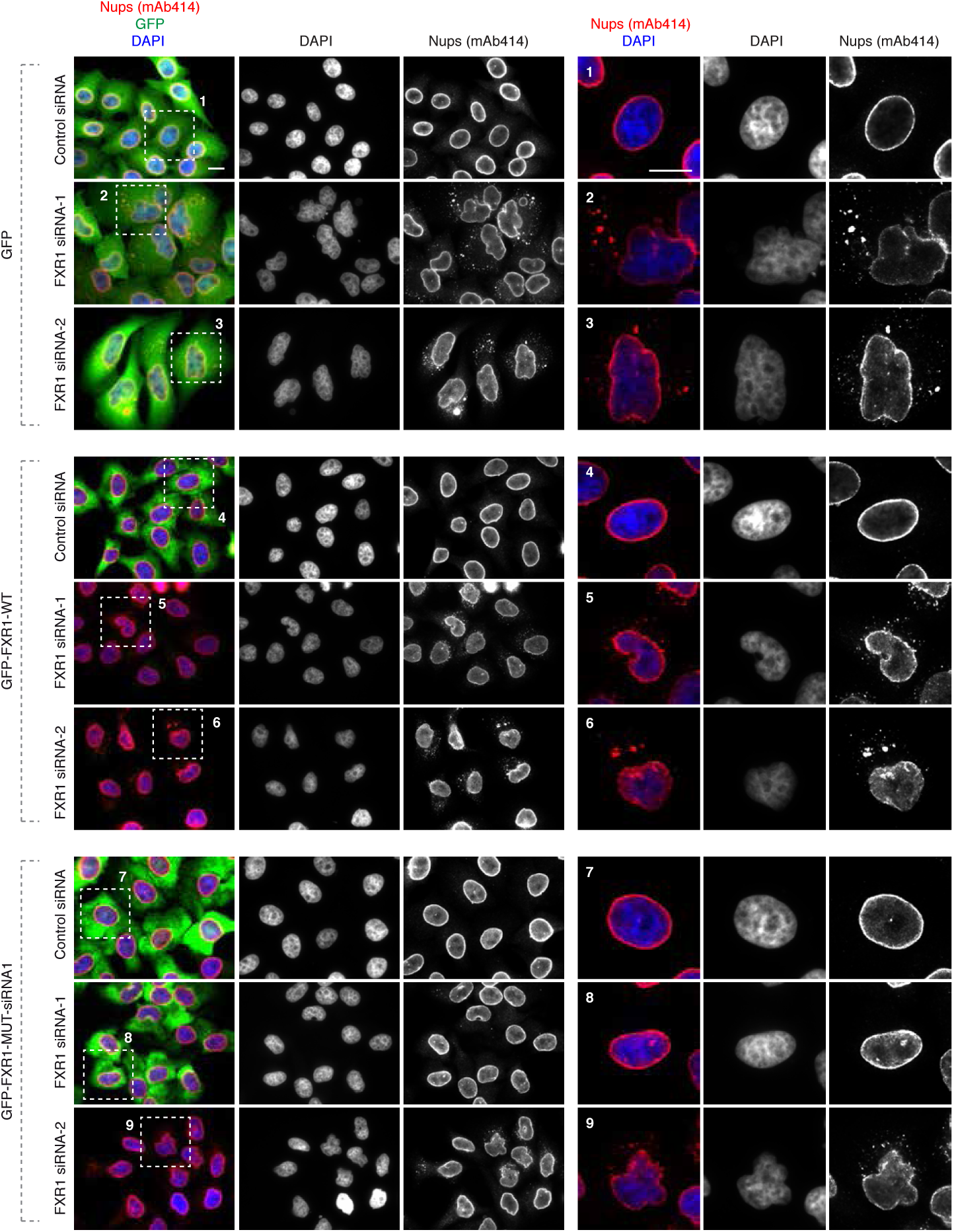
FXR1 specifically controls cytoplasmic Nups and nuclear shape. HeLa cells stably expressing GFP, GFP-FXR1 wild type (WT) and GFP-FXR1 mutated in the sequence recognized by FXR1 siRNA-1 (GFP-FXR1-MUT-siRNA1) were treated with the indicated siRNAs, synchronized by double thymidine block and released for 24 hours and analysed by immunofluorescence microscopy for mAb414. Depicted representative pictures correspond to the Western blot analysis shown in Figure 6B, quantifications of percentage of cells with cytoplasmic nucleoporin aggregates shown in Figure 2B and quantifications of percentage of cells with irregular nuclei shown in Figure 6C. Bars are 5 µm.

**Supplemental Figure 2:**
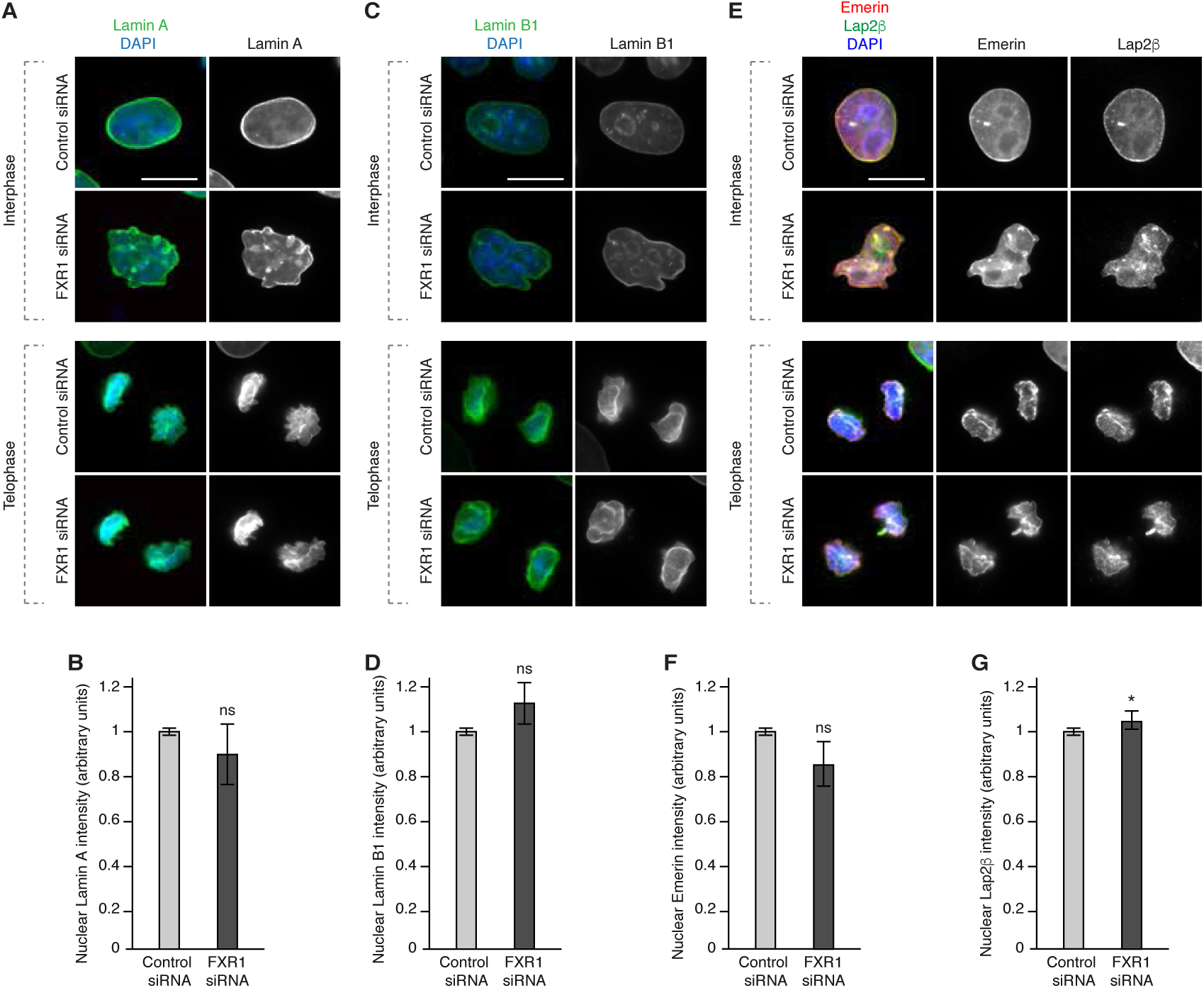
FXR1 does not drive recruitment of lamina-associated proteins. **(A-G)** HeLa cells were treated with the indicated siRNAs, synchronized by double thymidine block and released for 9 (telophase) and 12 (interphase) hours and analysed by immunofluorescence microscopy. The nuclear intensity of Lamin A (B), Lamin B1 (D), Emerin **(E)** and Lap2β (G) was quantified. ns indicates non-significant. Bars are 5 µm.

**Supplemental Figure 3:**
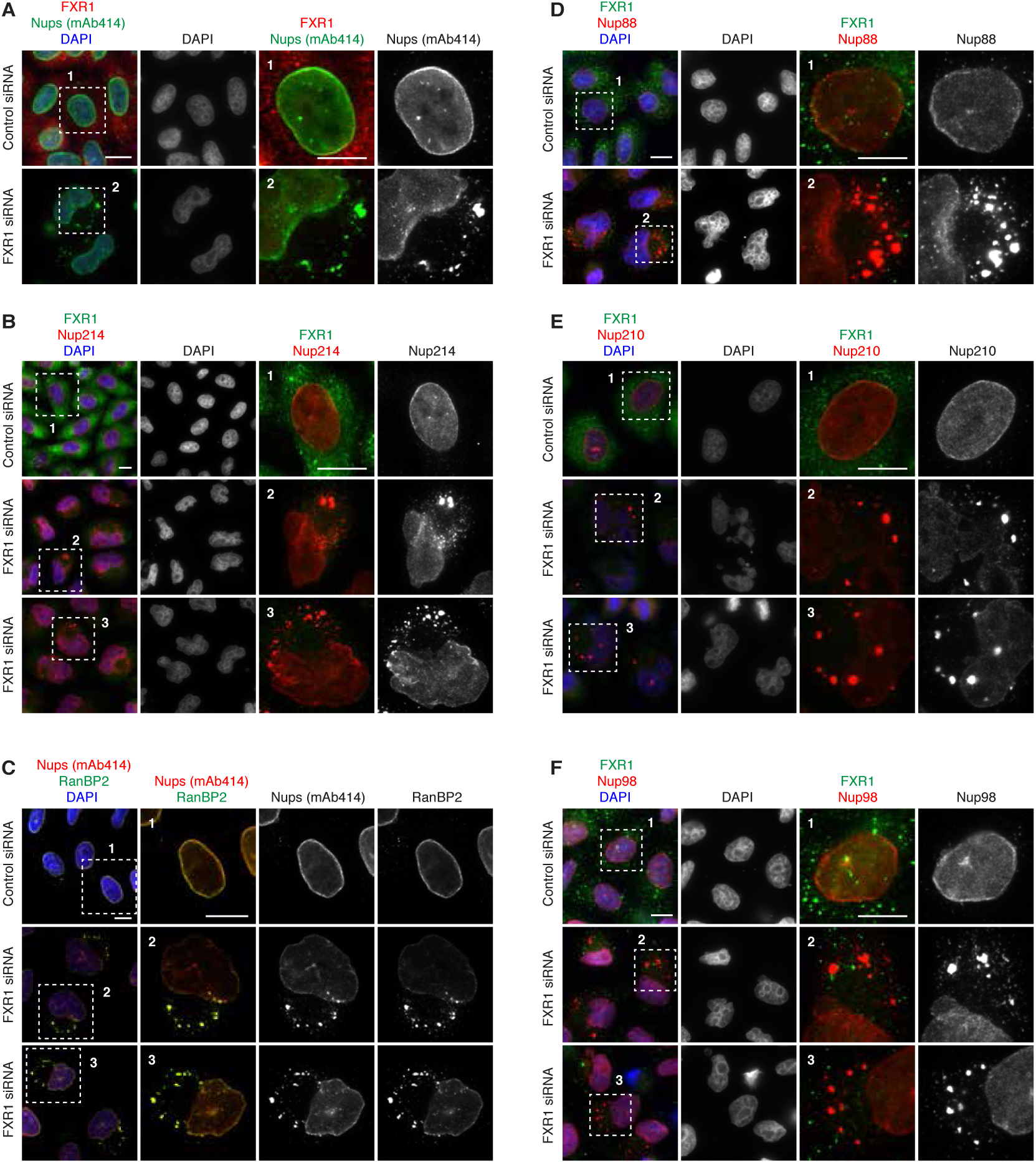
FXR1 inhibits aberrant assembly of cytoplasmic Nups. **(A-F)** HeLa cells were treated with the indicated siRNAs, synchronized by double thymidine block and released for 12 hours and analysed by immunofluorescence microscopy for indicated nucleoporins. The magnified framed regions are shown in the corresponding numbered panels. Additional or complementary representative images and channels of cells are shown, which are also depicted in Figure 2C. Bars are 5 µm.

**Supplemental Figure 4:**
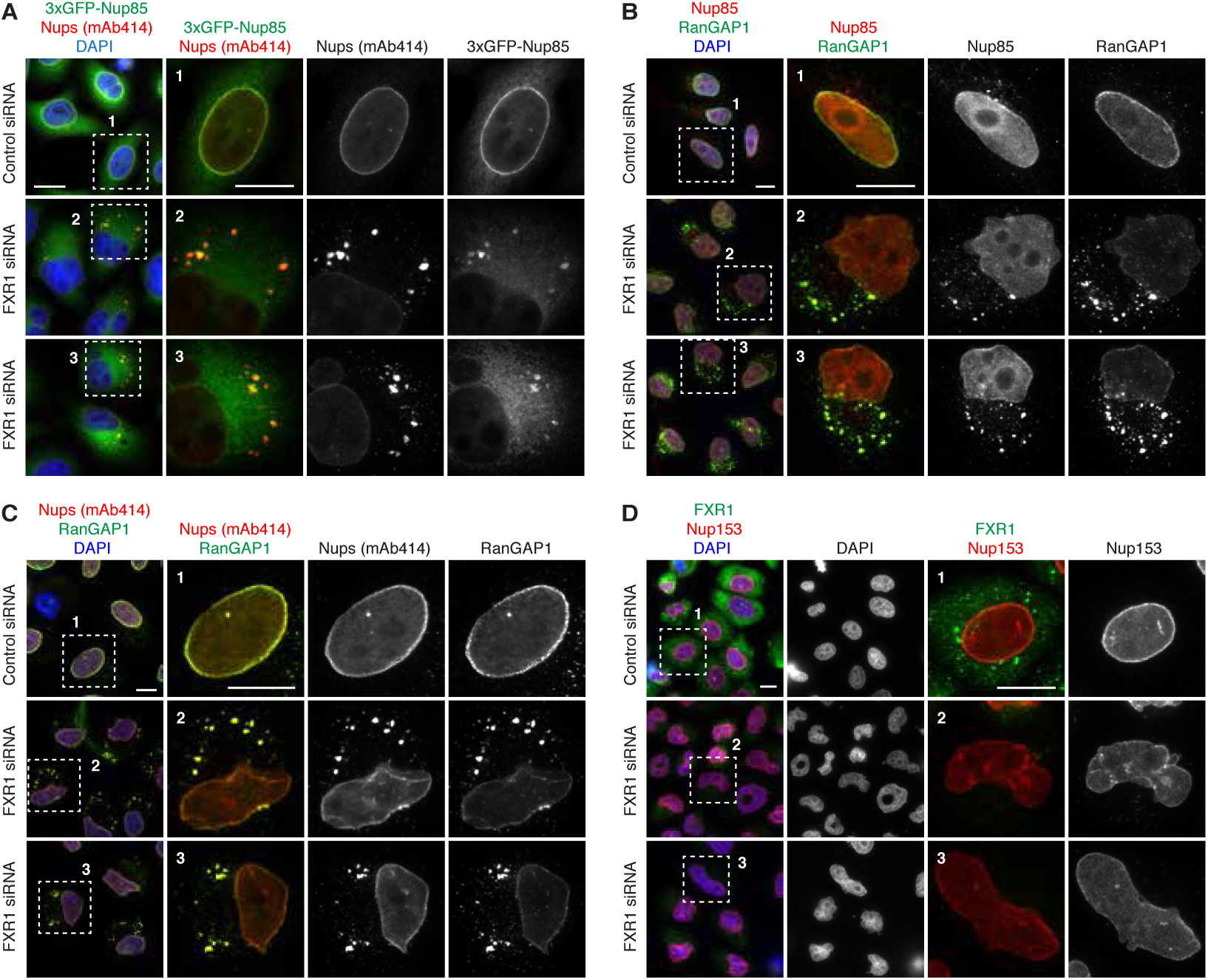
FXR1 inhibits aberrant assembly of cytoplasmic Nups. **(A-D)** HeLa cells stably expressing 3xGFP-Nup85 (A) or HeLa cells (B-D) were treated with the indicated siRNAs, synchronized by double thymidine block and released for 12 hours and analysed by immunofluorescence microscopy for indicated nucleoporins. The magnified framed regions are shown in the corresponding numbered panels. Additional or complementary representative images and channels of cells are shown, which are also depicted in Figure 2C. Bars are 5 µm.

**Supplemental Figure 5:**
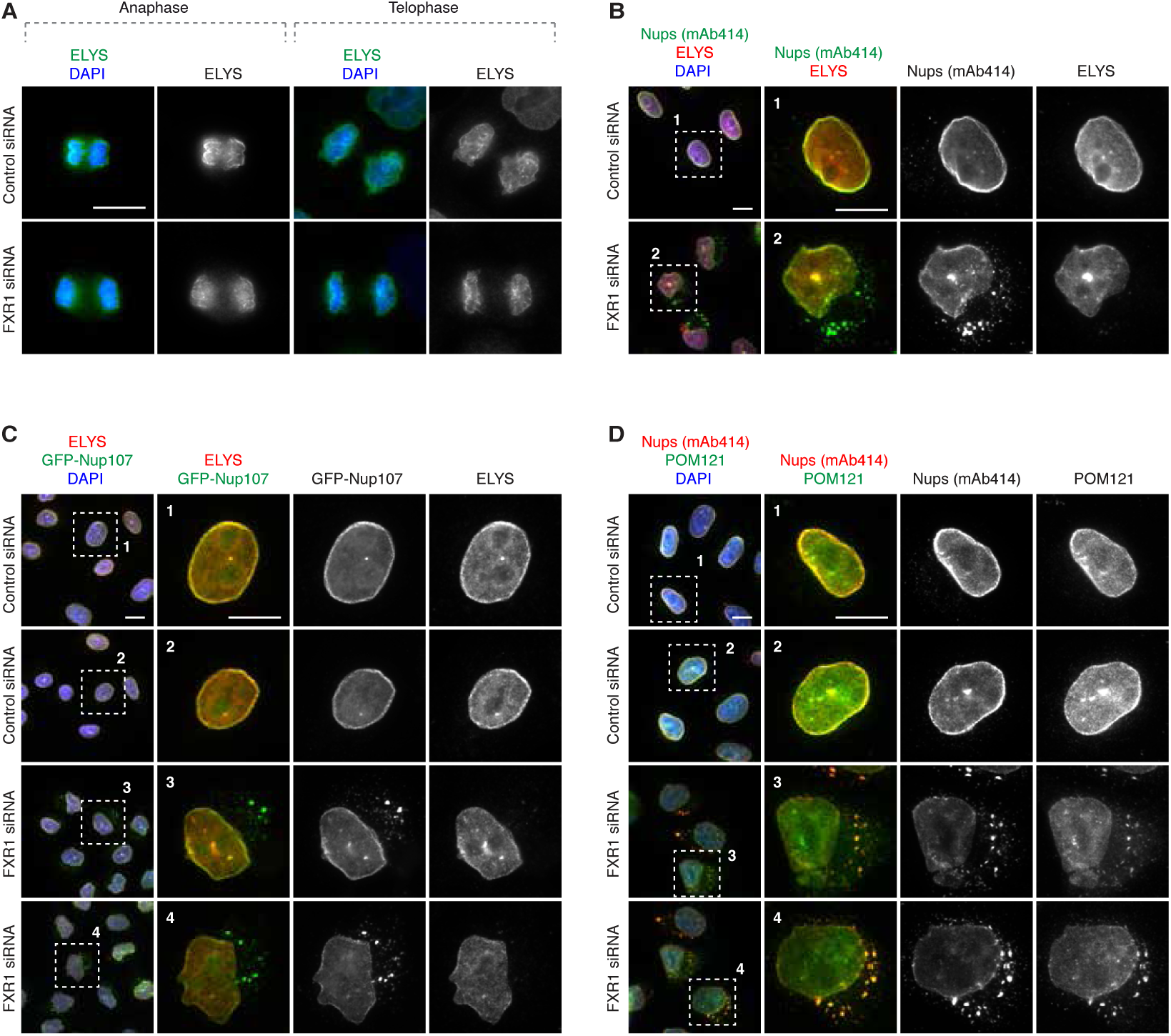
FXR1 inhibits aberrant assembly of cytoplasmic POM121 but not ELYS. **(A-D)** HeLa cells (A, B, D) or HeLa cells stably expressing GFP-Nup107 (C) were treated with the indicated siRNAs, synchronized by double thymidine block and released for 9 (telophase) and 12 (interphase) hours and analysed by immunofluorescence microscopy for ELYS and POM121. The magnified framed regions are shown in the corresponding numbered panels. Additional or complementary representative images and channels of cells are shown, which are also depicted in Figure 2C. Bars are 5 µm.

**Supplemental Figure 6:**
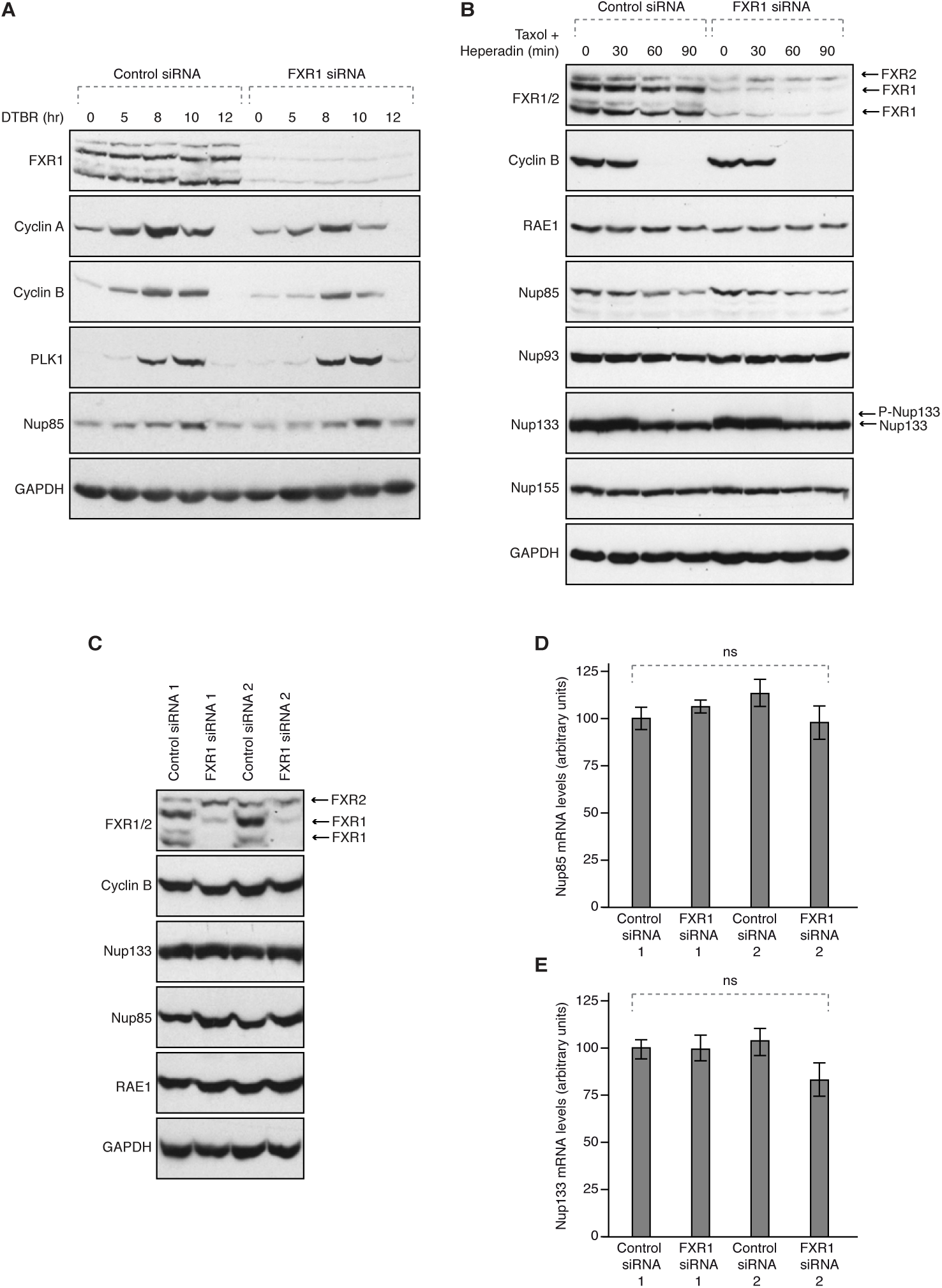
FXR1 does not regulate mitotic exit and protein levels of several Nups. **(A)** HeLa cells were treated with the indicated siRNAs, synchronized by double thymidine block and released for the indicated times (hours) and analysed by Western blotting. (**B)** HeLa cells were treated with the indicated siRNAs, synchronized by Hesperadin release from mitotic arrest induced by Taxol and analysed by Western blotting. Arrows point to FXR1 isoforms, FXR2 and phosphorylated (P-Nup133) and unmodified form of Nup133, respectively. (**C)** HeLa cells were treated with the indicated siRNAs and analysed by Western blotting. Arrows point to FXR1 isoforms and FXR2 protein. (**D-E)** HeLa cells were treated with the indicated siRNAs and mRNA levels of Nup85 (D) and Nup133 (E) were analysed by Quantitative Real-Time PCR. ns indicates non-significant.

**Supplemental Figure 7:**
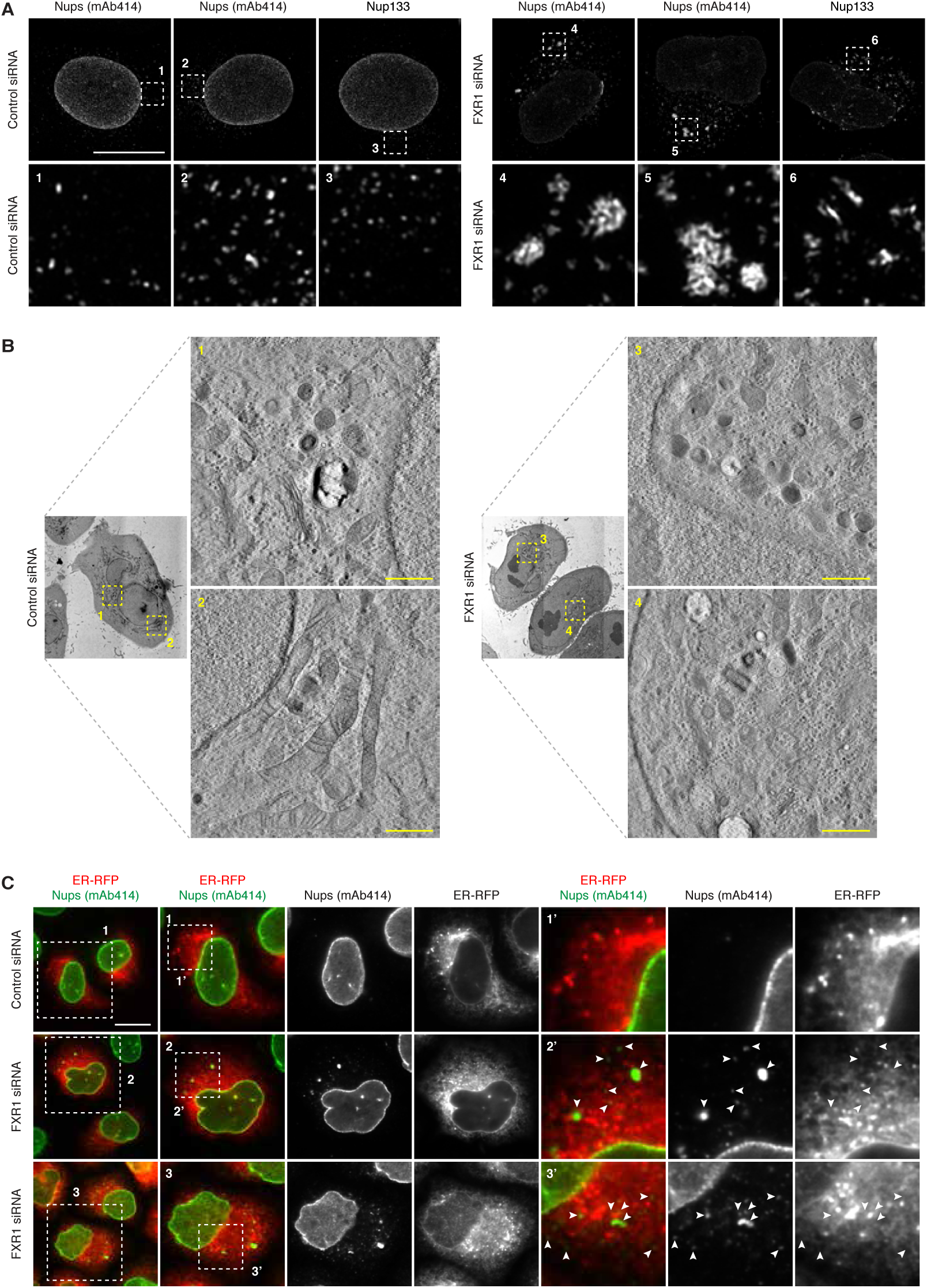
The cytoplasmic nucleoporin aggregates are Nup condensates distinct from ALs. **(A)** HeLa cells were treated with the indicated siRNAs, synchronized by double thymidine block and released for 12 hours and analysed by superresolution confocal microscopy. Bar is 5 µm. **(B)** HeLa cells were treated with the indicated siRNAs, synchronized by double thymidine block and released for 12 hours and analysed by electron microscopy. The magnified framed regions are shown in the corresponding numbered panels. Bar is 1 µm. **(C)** HeLa cells were treated with indicated siRNAs, transfected with ER-RFP reporter for 24h, synchronized by double thymidine block and released for 12 hours and analysed by immunofluorescence microscopy for mAb414. The magnified framed regions are shown in the corresponding numbered panels. Arrowheads point to the FG-Nup-positive cytoplasmic aggregates. Bar is 5 µm.

**Supplemental Figure 8:**
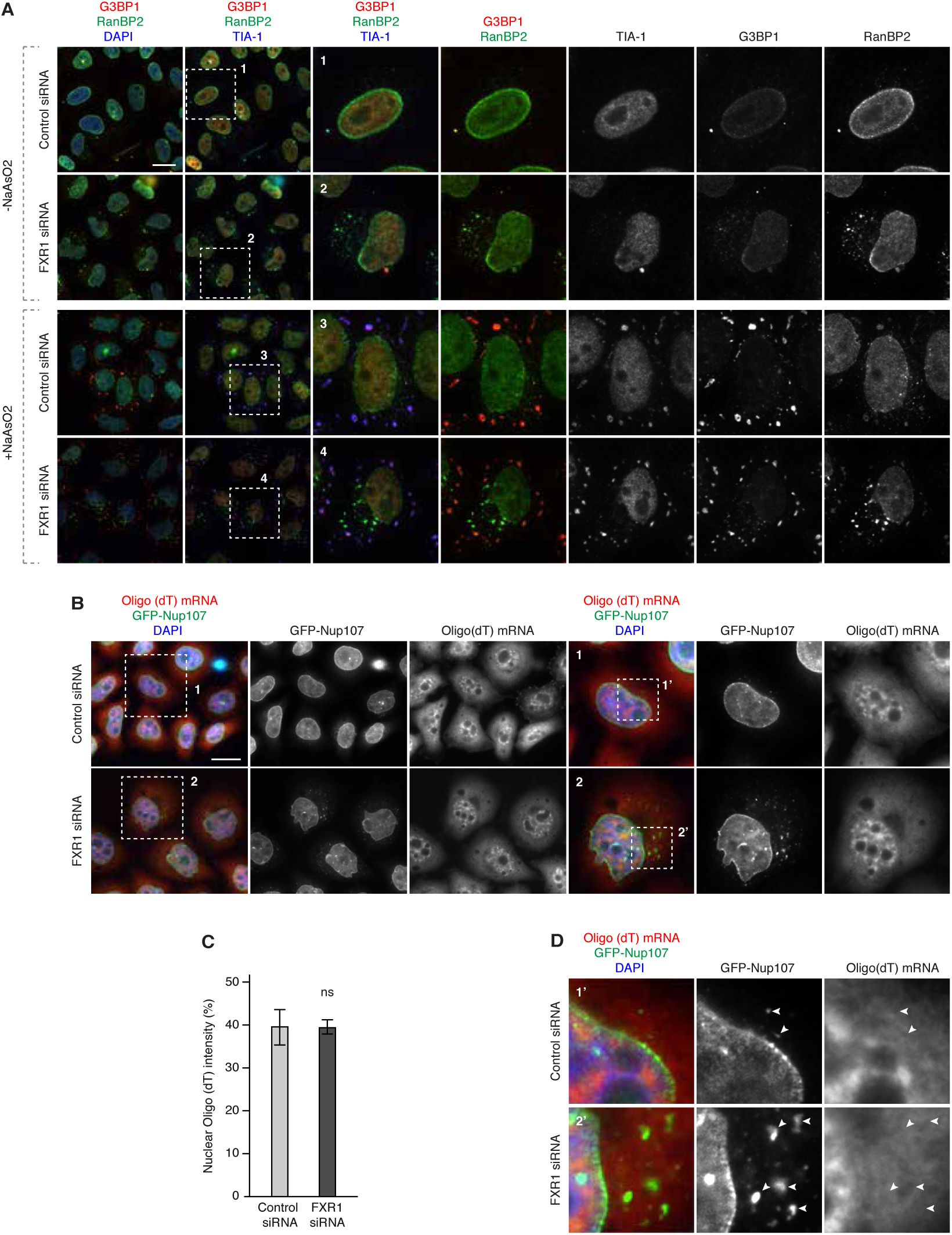
The cytoplasmic nucleoporin aggregates are distinct from SGs and do not contain mRNAs. **(A)** HeLa cells were treated with the indicated siRNAs, synchronized by double thymidine block and released for 12 hours, treated with or without the stress inducing factor NaAsO2 0,5 mM for 1 hour and analysed by immunofluorescence microscopy for the stress granule markers G3BP1 and TIA-1 and the nucleoporin RanBP2. The magnified framed regions are shown in the corresponding numbered panels. **(B-D)** HeLa cells stably expressing GFP-Nup107 were treated with the indicated siRNAs, synchronized by double thymidine block and released for 12 hours, hybridized with the oligo d(T) mRNA FISH probe and analysed by immunofluorescence microscopy. The magnified framed regions are shown in the corresponding numbered panels in (B) and higher magnifications are shown in (D). Arrowheads point to GFP-Nup107-positive cytoplasmic aggregates. The percentage of nuclear mRNA intensity (n=600) was quantified in (C). ns indicates non-significant. Bars are 5 µm.

**Supplemental Figure 9:**
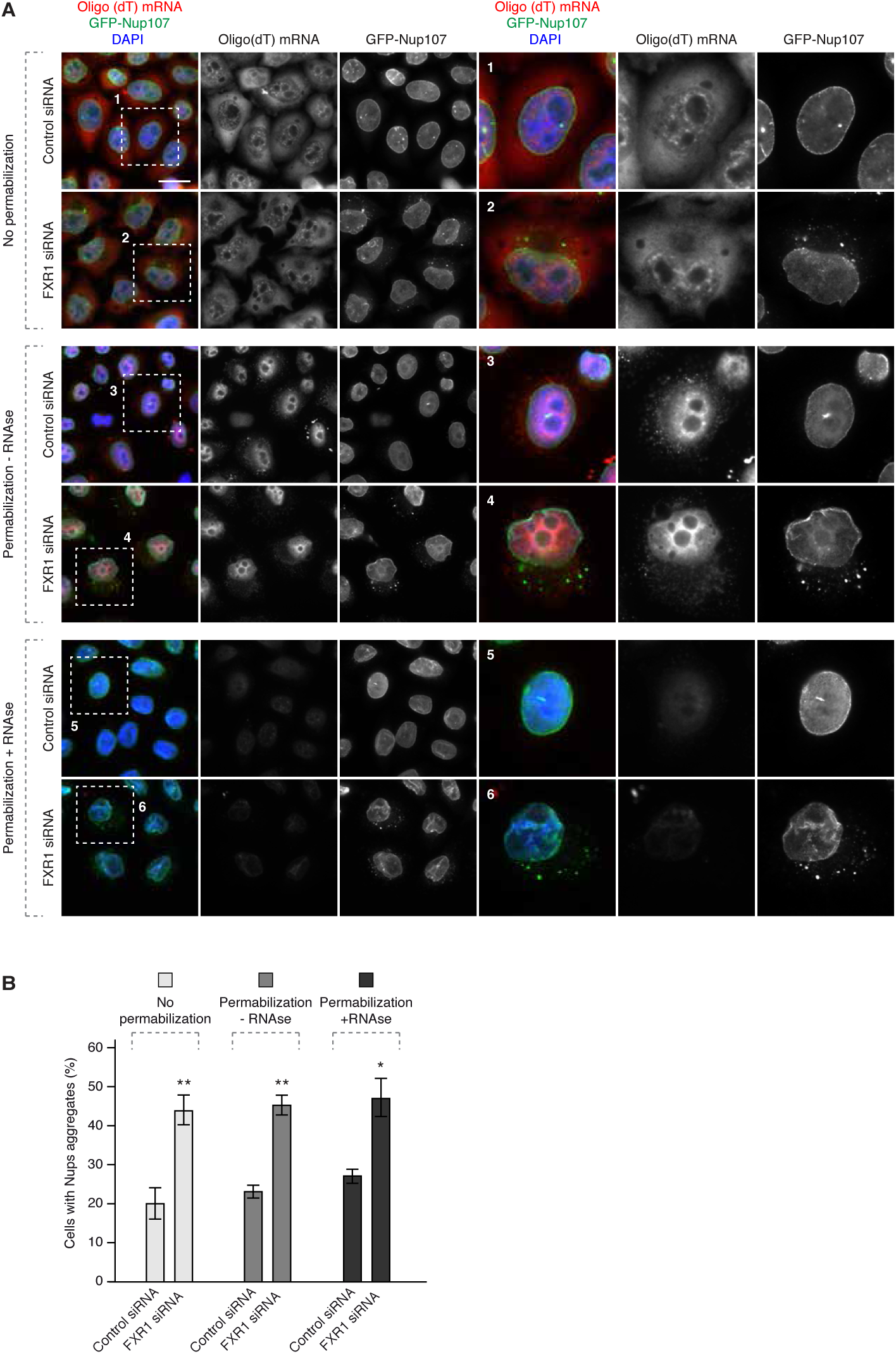
RNAs are dispensable for maintenance and dynamics of the cytoplasmic nucleoporin aggregates. **(A-B)** HeLa cells stably expressing GFP-Nup107 were treated with the indicated siRNAs, synchronized by double thymidine block and released for 12 hours, hybridized with the oligo d(T) mRNA FISH probe, treated with or without RNaseA/T1 and/or digitonin permabilization for 5 min and analysed by fluorescence microscopy. The magnified framed regions are shown in the corresponding numbered panels. The percentage of cells with GFP-Nup107-positive cytoplasmic aggregates (n=8900) was quantified in (B). Bar is 5 µm.

**Supplemental Figure 10:**
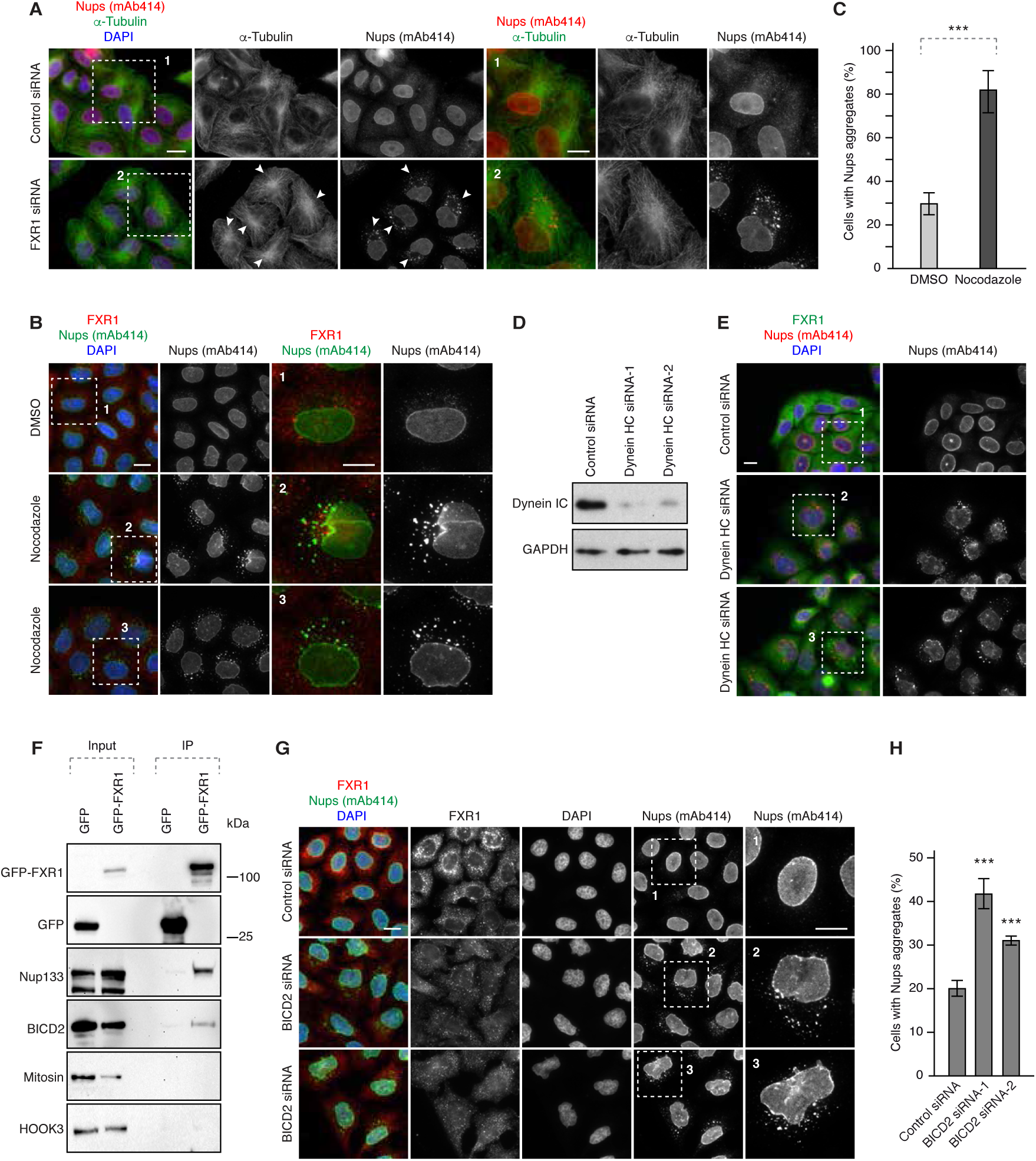
FXR1 inhibits formation of the cytoplasmic nucleoporin aggregates by dynein-BICD2-based microtubule-dependent transport. **(A)** HeLa cells were treated with the indicated siRNAs, synchronized by double thymidine block and released for 12 hours and analysed by immunofluorescence microscopy for α-tubulin and mAb414. Arrowheads point to the positions of the cytoplasmic nucleoporin aggregates relative to MTOC. **(B-C)** HeLa cells were synchronized by double thymidine block and released for 12 hours, treated with nocodazole or solvent (DMSO) for 90 min and analysed by immunofluorescence microscopy for FXR1 and mAb414. The magnified framed regions are shown in the corresponding numbered panels in (B). The percentage of cells with cytoplasmic nucleoporin aggregates (n=2500) was quantified in (C). **(D-E)** HeLa cells were treated with the indicated siRNAs, synchronized by double thymidine block and released for 12 hours and analysed by Western blotting and immunofluorescence microscopy. The magnified framed regions of pictures shown in (E) are depicted in the corresponding numbered panels in the Figure 4B. **(F)** Lysates of HeLa cells stably expressing GFP alone or GFP-FXR1 were subjected to immunoprecipitation using GFP-Trap beads (GFP-IP) and analysed by Western blotting. (**G-H)** HeLa cells were treated with the indicated siRNAs, synchronized by double thymidine block and released for 12 hours and analysed by immunofluorescence microscopy for FXR1 and mAb414. The magnified framed regions are shown in the corresponding numbered panels in (G). The percentage of cells with cytoplasmic nucleoporin aggregates (n=3500) was quantified in (H). Bars are 5 µm.

**Supplemental Figure 11:**
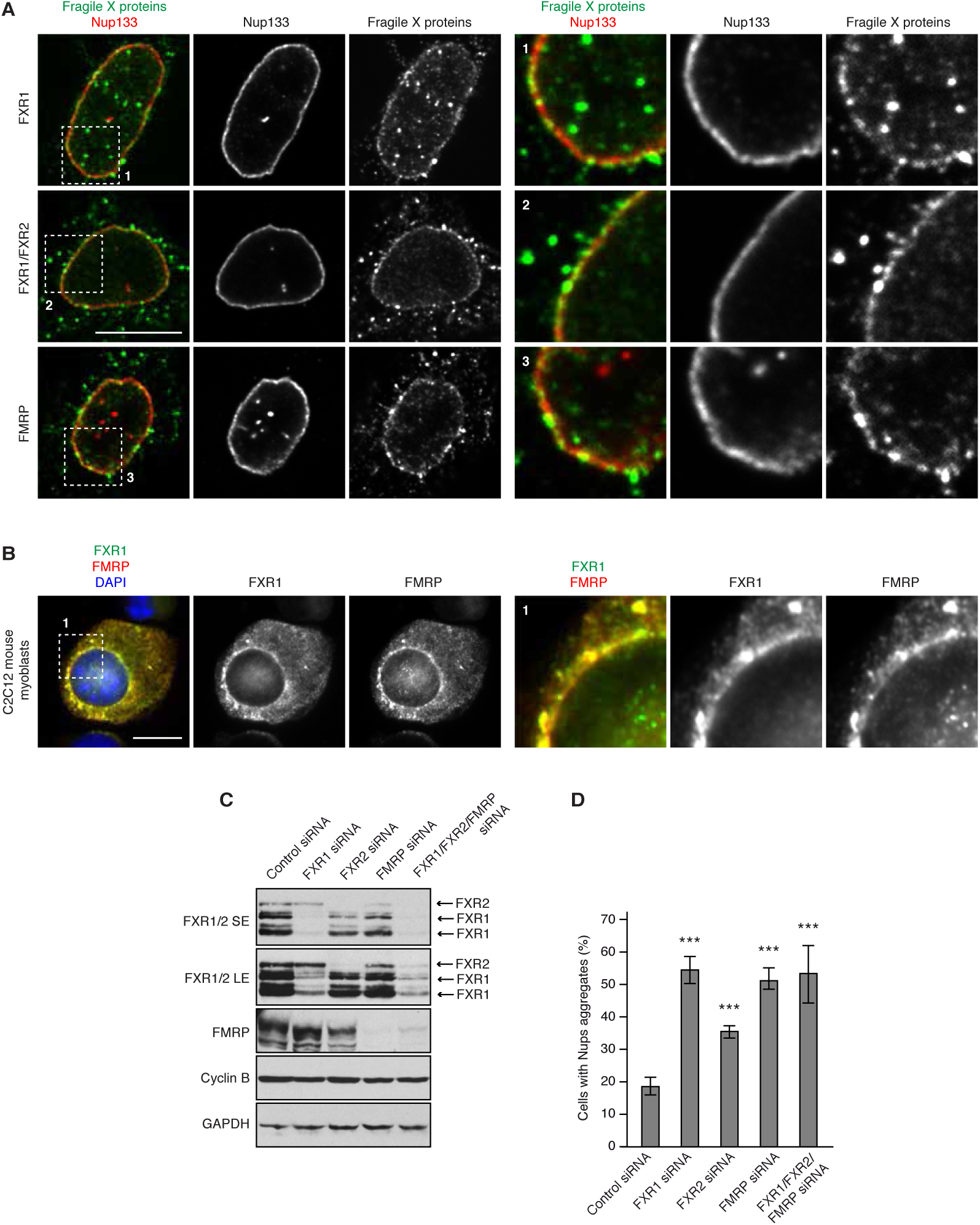
FXR protein family members localize to NE and regulate cytoplasmic Nups. **(A-B)** HeLa cells (A) and mouse myoblasts (B) were analysed by immunofluorescence microscopy for FXR1, FXR1+2 and FMRP. The magnified framed regions are shown in the corresponding numbered panels. Bars are 5 µm. **(C-D)** HeLa cells were treated with the indicated siRNAs, synchronized by double thymidine block and released for 12 hours and analysed by Western blotting and immunofluorescence microscopy. The percentage of cells with cytoplasmic nucleoporin aggregates (n=900) was quantified in (D).

**Supplemental Figure 12:**
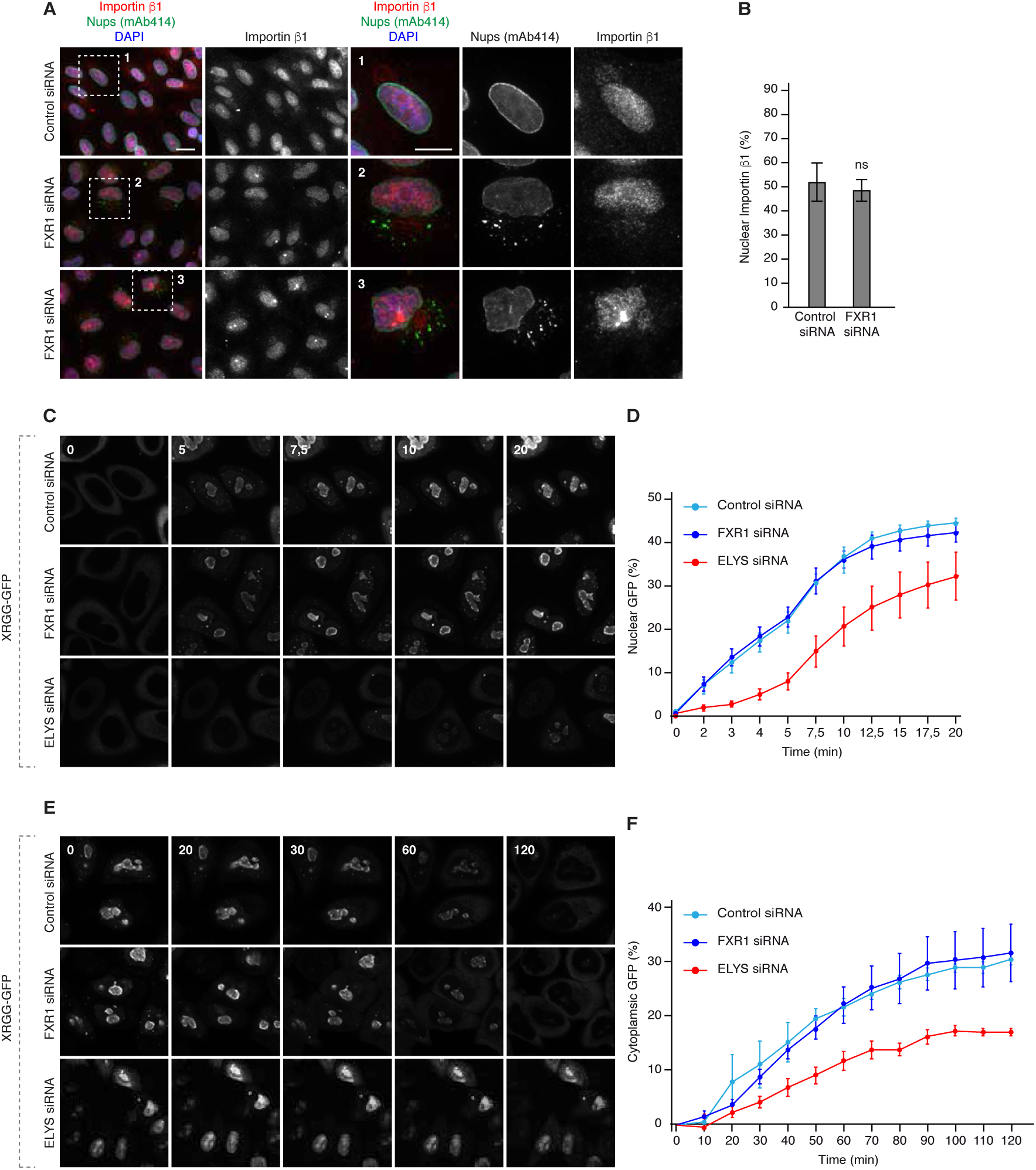
FXR1 protein does not regulate general nucleocytoplasmic transport. **(A-B)** HeLa cells were treated with the indicated siRNAs, synchronized by double thymidine block and released for 12 hours and analysed by immunofluorescence microscopy. The percentage of nuclear Importin β1 intensity (n=300) was quantified in (B). **(C-D)** HeLa cells were transfected with the import reporter plasmid XRGG-GFP, treated with the indicated siRNAs and synchronized in early G1 phase by Monastrol release. Dexamethasone-induced nuclear import of XRGG-GFP was analysed by live video spinning disk confocal microscopy. The selected frames of the movies are depicted and time is shown in minutes in (C). The percentage of nuclear XRGG-GFP over time (n=247) was quantified in (D). **(E-F)** HeLa cells were treated as in (C-D) and Dexamethasone was added for 3 hours to induce XRGG-GFP nuclear import. Following wash-out the nuclear export of XRGG-GFP was analysed by live video spinning disk confocal microscopy. The selected frames of the movies are depicted and time is shown in minutes in (E). The percentage of cytoplasmic XRGG-GFP over time (n=174) was quantified in (F).

